# Multi-dimensional regulation of LIN-28 temporal expression dynamics in the *C. elegans* heterochronic gene cascade

**DOI:** 10.1101/2025.11.13.688268

**Authors:** Charles Nelson, Victor Ambros

**Author notes:** Corresponding author: Charles Nelson, Program in Molecular Medicine, University of Massachusetts Chan Medical School, Worcester, MA 01605, USA.

## Abstract

LIN-28 is an evolutionarily conserved RNA-binding protein that plays critical roles in regulating pluripotency and cell fate determination during animal development. In *Caenorhabditis elegans*, *lin-28* is an integral component of the heterochronic (developmental timing) gene regulatory cascade. Loss-of-function mutations in *lin-28* result in precocious cell fate determination during larval development. Previous studies showed that the proper progression of stage specific cell fates during larval development depends on the progressive downregulation of LIN-28, which is negatively regulated by the *lin-4* microRNA through complementary sequences located in the *lin-28* 3’ UTR. In this study, we employ CRISPR/Cas9 editing of the endogenous *lin-28* locus to demonstrate that the robust developmental downregulation of LIN-28 involves contributions from multiple inputs. These include a convergent action of the *let-7* family and *lin-4* microRNAs via adjacent complementary sites in the *lin-28* 3’ UTR, in conjunction with the previously described post-translational inhibition of LIN-28 by the *lep-5* long non-coding RNA, which all together account for virtually the entirety of LIN-28 repression. Furthermore, through the systematic testing of a series of truncations of the *lin-28* 3’ UTR, we identify three positive regulatory regions that enhance LIN-28 expression, thereby counterbalancing the negative effects of the *let-7* and *lin-4* microRNAs and the *lep-5* long non-coding RNA.

## INTRODUCTION

To achieve a properly developed animal, the precise orchestration of gene expression is essential for establishing correct cell lineages and cell fates at appropriate developmental stages. *Caenorhabditis elegans* is a hermaphroditic multicellular animal that undergoes six distinct developmental stages—embryogenesis, the L1-L4 larval stages, and the adult stage—and exhibits invariant spatial and temporal patterns of cell lineage fates, resulting in an adult with 959 somatic nuclei (Sulston and Horvitz 1977). In *C. elegans*, the temporally dynamic heterochronic (developmental timing) gene regulatory cascade governs the precise timing of cell-fate transitions during larval development via a hierarchical arrangement of genes that either restrict or promote these transitions (Rougvie and Moss 2013). Central to the heterochronic cascade are the transcription factors LIN-14 and HBL-1, as well as the RNA binding protein LIN-28. Loss-of-function mutations in *lin-14*, *lin-28*, or *hbl-1* result in the skipping of early larval cell fate transitions, leading to precocious expression of later cell fate programs, whereas gain-of-function alleles cause the reiteration of early larval cell fates, resulting in so-called retarded development (Ambros and Horvitz 1984; Fay et al. 1999; Ilbay and Ambros 2019).

MicroRNAs (miRNAs) are small, non-coding RNAs that play a crucial role in the negative regulation of protein expression, primarily through the base-pairing of nucleotides 2-8 of the miRNA—referred to as the “seed” sequence—to complementary sequences located within the 3’ untranslated regions (UTRs) of target messenger RNAs (mRNAs). miRNAs that possess an identical seed sequence but differ in their non-seed nucleotides can target the same mRNAs and are classified within the same miRNA family. While seed-only base-pairing can suffice to inhibit target expression, the involvement of non-seed pairing between the miRNA and its target mRNA can enhance both the potency and specificity of gene repression (Bartel 2009; Brancati and Grosshans 2018; Ambros and Ruvkun 2018; Duan et al. 2022).

In *C. elegans*, the microRNAs miR-48, miR-84, miR-241, and *let-7* microRNA (hereafter collectively referred to as *let-7* family microRNAs unless specified otherwise) and the *lin-4* microRNA facilitate transitions from early to later larval developmental events. Loss-of-function mutations of genes encoding the *lin-4* microRNA or the *let-7* family microRNAs lead to retarded heterochronic phenotypes (Chalfie et al. 1981; Reinhart et al. 2000; Abbott et al. 2005). The 3’ UTR regions of the genes *lin-14*, *lin-28*, and *hbl-1* contain sequences with full seed complementarity to the *lin-4* microRNA and/or the *let-7* family microRNAs (Lee et al. 1993; Moss et al. 1997; Reinhart et al. 2000; Abrahante et al. 2003; Lin et al. 2003). Hereafter, we refer to these microRNA complementary sequences as *lin-4*-*CSs* and *let-7-CSs* (for *<u>C</u>*omplementary *<u>S</u>*equence*s*). We note that herein the terms “*let-7-CS”* or *“let-7* microRNA” are intended to encompass potential regulation by any or all the *let-7* family microRNAs. Similarly, the terms “*lin-4-CS”* or *“lin-4* microRNA” encompass potential regulation by *lin-4* itself or the other member of the *lin-4* family, *mir-237*.

Deletions within the endogenous 3’ UTRs of either *lin-14* or *hbl-1* that eliminate *lin-4*-*CSs* and *let-7-CSs* result in pronounced gain-of-function phenotypes that phenocopy the loss of the *lin-4* or *let-7* family microRNAs (Wightman et al. 1993; Ilbay and Ambros 2019). The strikingly similar phenotypes associated with these extensive 3’ UTR deletions to those observed in the absence of the *lin-4* and/or *let-7* family microRNAs suggest that microRNA-mediated repression via the 3’ UTRs of *lin-14* and *hbl-1* constitutes the predominant regulatory mechanism for the downregulation of LIN-14 and HBL-1.

LIN-28 (Lin28 in mammals; hereafter referred to as LIN-28 unless otherwise noted) is an evolutionarily conserved RNA-binding protein that interacts with and regulates a variety of mRNAs and noncoding RNAs (Wilbert et al. 2012; Stefani et al. 2015; Kiontke et al. 2019). In animals ranging from *C. elegans* to humans, LIN-28 can bind to the primary and precursor transcripts of the *let-7* microRNA to inhibit its function through various mechanisms, including the blockage of transcript processing, transcript degradation, and the promotion of primary transcript trans-splicing (Rybak et al. 2008; Viswanathan et al. 2008; Stefani et al. 2015; Nelson and Ambros 2019). The loss of LIN-28 can lead to the overexpression of the *let-7* microRNA, resulting in premature cellular differentiation and development (Johnson et al. 2003; Vadla et al. 2012). In wild-type *C. elegans*, LIN-28 expression is highest during the L1 and L2 stages and significantly diminishes by the L3 stage (Seggerson et al. 2002). Conversely, the expression of the *lin-4* microRNA increases during mid to late L1 larvae, while the levels of *let-7* microRNAs rise during the L2-L4 stages (Feinbaum and Ambros 1999; Reinhart et al. 2000; Abbott et al. 2005; Nelson and Ambros 2021).

The *C. elegans lin-28* 3’ UTR contains a *let-7*-*CS* and a *lin-4-CS* with full seed complementarity to their respective microRNAs, and therefore, it is posited that the *lin-4* microRNA and the *let-7* microRNAs inhibit LIN-28 accumulation during later larval stages (Moss et al. 1997; Reinhart et al. 2000). Currently, no mutations have been identified in the endogenous *C. elegans lin-28* 3’ UTR, leaving the relative contributions of the *lin-4* and *let-7* microRNAs to LIN-28 down regulation undetermined. The expression of a high copy *lin-28* transgene with the *lin-4-CS* deleted resulted in misexpression of LIN-28 from the transgene and led to pronounced gain-of-function phenotypes (Moss et al. 1997). However, the potential confounding effects associated with the high copy nature of such transgenes introduce uncertainly whether these previous results accurately represent the endogenous regulation of LIN-28 expression.

MicroRNAs are known to repress target gene expression by regulating mRNA levels and/or translational efficiency. Therefore, expression of the *lin-4* and *let-7* microRNAs contribute to LIN-28 down regulation by inhibiting LIN-28 production. An additional layer of repression of LIN-28 is exerted by the *lep-5* long non-coding RNA (lncRNA), which is believed to destabilize LIN-28 by binding to LIN-28 and recruiting the E3 ubiquitin-protein ligase LEP-2 (Herrera et al. 2016; Kiontke et al. 2019). Expression of the *lep-5* lncRNA begins to increase in the L2 stage and is hypothesized to eliminate residual LIN-28 that was translated during embryogenesis and the early L1 stage, prior to the expression of the *lin-4* and *let-7* microRNAs (Kiontke et al. 2019). Importantly, the loss of *lep-5* does not fully derepress LIN-28, which is consistent with the developmental down regulation of LIN-28 reflecting contributions from both microRNA-mediated repression and *lep-5* lncRNA-induced destabilization of LIN-28 (Kiontke et al. 2019).

Here, we show that the *let-7*-*CS* and the *lin-4*-*CS* targeting of LIN-28 mRNA is conserved amongst *Caenorhabditis* species. Furthermore, we demonstrate that the *let-7-CS* and the *lin-4-CS* located within the 3’ UTR of *C. elegans lin-28* function semi-redundantly to downregulate LIN-28 production during larval development. Our data also suggest that this microRNA-mediated downregulation of LIN-28 production acts synergistically with the post-translational degradation of LIN-28 mediated by the *lep-5* lncRNA. Remarkably, we observed that the deletion of the entire *lin-28* 3’ UTR results in a significantly less severe phenotype than deletion of only the CSs. This led us to identify three positive regulatory regions within the *lin-28* 3’ UTR that promote LIN-28 expression by stabilizing its mRNA. Our data further suggests that the various modes of regulation of LIN-28 can be conditional: 3’ UTR mutants can exhibit exacerbated heterochronic phenotypes in response to further alterations in LIN-28 expression or in response to environmental stresses, including exposure to the pathogenic bacterium *Pseudomonas aeruginosa* or development at various temperatures.

Our findings illustrate that multiple regulatory layers govern LIN-28 developmental dynamics. These include microRNAs and positive elements acting via the 3’ UTR, which affect LIN-28 mRNA stability and LIN-28 protein synthesis, and the *lep-5* lncRNA, which functions independently of LIN-28 mRNA to regulate LIN-28 protein stability. We propose that these layered regulatory mechanisms are crucial for ensuring precise expression and timely downregulation of LIN-28, irrespective of environmental conditions.

## RESULTS

### The targeting of LIN-28 mRNA by the *let-7* and *lin-4* microRNAs is conserved among *Caenorhabditis* species

The *C. elegans lin-28* 3’ UTR contains one *let-7-CS* and one *lin-4-CS*, both of which exhibit full seed complementarity to their respective microRNAs and are situated 13 nucleotides between seed pairings (Fig. 1 and Table S1). To assess the conservation of *let-7* and *lin-4* microRNA targeting of LIN-28 mRNA across various *Caenorhabditis* species, we identified homologs of *C. elegans lin-28* using BLAST searches based on the coding sequence and subsequently examined the downstream sequence (likely comprising 3’ UTR regions) for the presence of *let-7-CSs* and *lin-4-CSs* followed by a polyadenylation sequence (PAS). All species analyzed possessed a single copy of *lin-28*, with the exception of *C. brenneri* and *C. japonica*, both of which contained two copies (Fig. 1 and Table S1). Furthermore, all species exhibited probable PASs of AATAAA located between 172 and 1544 base pairs from the stop codon, except for *C. japonica*, which displayed likely PASs of AATAAG for both paralogs (Table S1).

**Figure 1.**
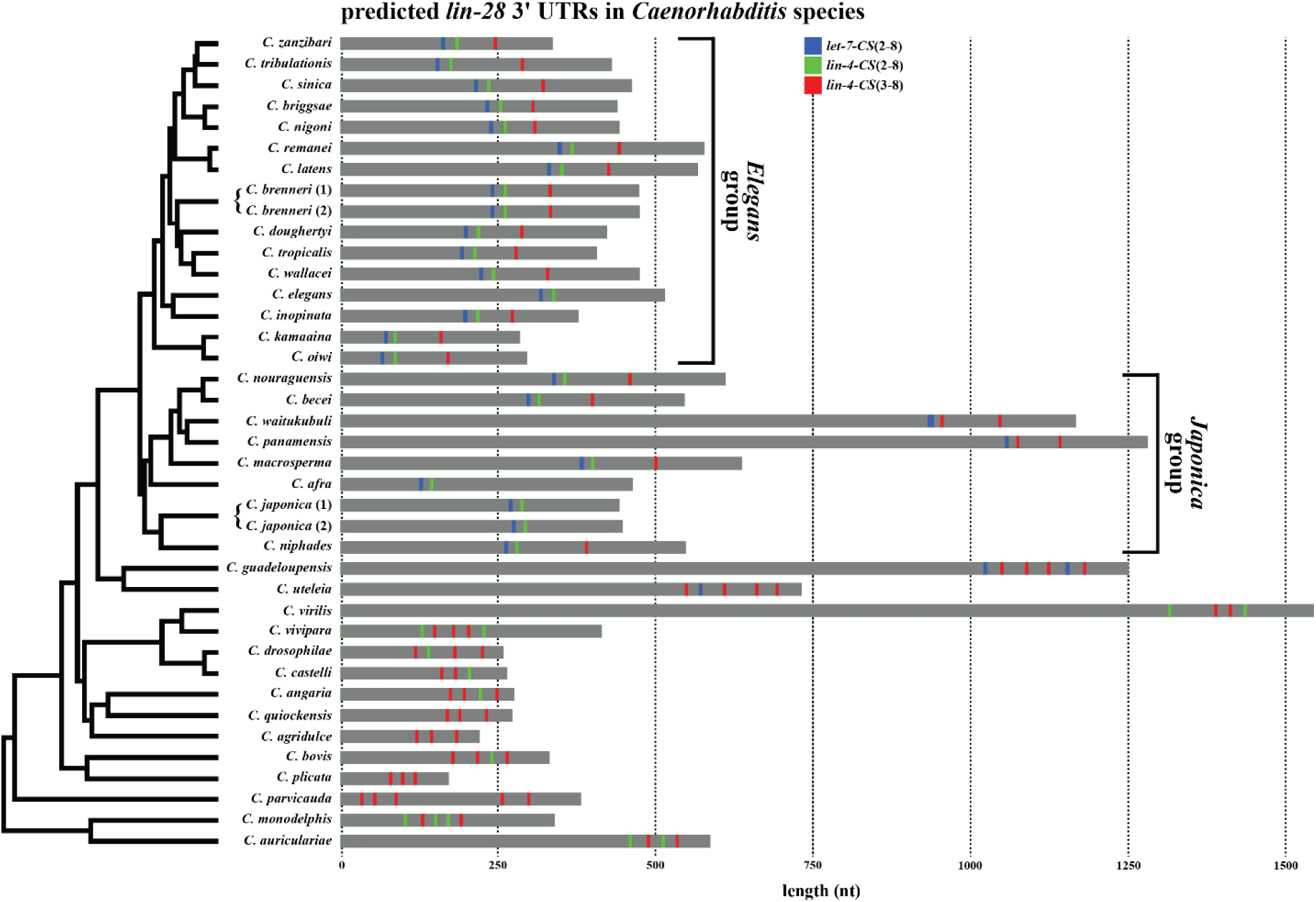
Conservation of the *let-7* and *lin-4* microRNA-binding sites in the *lin-28* 3’ UTRs of *Caenorhabditis* species. Predicted *let-7*-*CSs* and *lin-4-CSs* in the probable *lin-28* 3’ UTRs of *Caenorhabditis* species. Blue and green boxes denote the respective locations of *let-7*-*CSs* and *lin-4*-*CSs* that exhibit predicted base pairing with nucleotides two through eight of their corresponding microRNAs. Red boxes indicate the locations of *lin-4*-*CSs* predicted to base pair with nucleotides three through eight of the *lin-4* microRNA. Phylogeny is adapted from previous publications (Sun et al. 2022; Picao-Osorio et al. 2025).

Similar to *C. elegans*, all predicted *lin-28* 3’ UTR regions in the *Elegans* group of *Caenorhabditis* species contained a *let-7-CS* with complementarity to nucleotides 2-8 (hereafter referred to as *let-7*-*CS(2-8)*, with (2-8) representing the complementary nucleotides of the microRNA’s seed; Fig. 1 and Table S1). In all species within the *Elegans* group, the *let-7-CS* was followed by a *lin-4*-*CS(2-8)* (with an average distance of 13 ± 1.5 nt between seed pairings; Fig. 1 and Table S1). Notably, we also found that all 3’ UTR regions in the *Elegans* group, with the exception of *C. elegans*, harbored a second *lin-4*-*CS* with complementarity to nucleotides 3-8 of the *lin-4* microRNA (hereafter referred to as *lin-4*-*CS(3-8)*) located further downstream in the predicted 3’ UTRs (Fig. 1 and Table S1). Similarly, in the *Japonica* group, all *lin-28* 3’ UTR regions exhibited a *let-7-CS(2-8)* that was closely followed by a *lin-4-CS(2-8)* (present in seven of nine species) or a *lin-4-CS(3-8)* (present in two of nine species) (Fig. 1 and Table S1). Furthermore, in the *Japonica* group, six of the nine 3’ UTR regions also contained an additional *lin-4-CS(3-8)* located downstream of the first *lin-4-CS* (Fig. 1 and Table S1). Outside of the *Elegans* and *Japonica* groups, only two species analyzed, *C. guadeloupensis* and *C. uteleia*, which are the species most closely related to the *Elegans* and *Japonica* groups, exhibited a *let-7-CS* in their *lin-28* 3’ UTR regions (Fig. 1 and Table S1). All remaining analyzed *Caenorhabditis lin-28* 3’ UTR regions exhibited between two and five *lin-4-CSs*, with seven species containing only *lin-4-CS(3-8)* and the remaining species harboring at least one *lin-4-CS(2-8)* along with multiple *lin-4-CS(3-8)* (Fig. 1 and Table S1).

### The *let-7* and *lin-4* microRNAs act redundantly to regulate the expression of LIN-28

MicroRNAs post-transcriptionally inhibit the expression of protein-coding genes (Shang et al. 2023). Accordingly, we hypothesized that the loss of the *let-7* and/or *lin-4* microRNA-mediated targeting of the LIN-28 mRNA in *C. elegans* would lead to retarded developmental phenotypes due to the misexpression (gain-of-function) of LIN-28 protein. In *C. elegans*, retarded developmental phenotypes frequently manifest as defects in vulval formation and integrity, as well as abnormal expression patterns of the adult-specific transgenic reporter *pCol-19::GFP* (Reinhart et al. 2000; Abbott et al. 2005). To quantify the penetrance of retarded development, we devised two scoring methods for young adult specimens: 1) assessing the penetrance of vulval dysfunction (referred to as “adult phenotype”; Fig. 2a), and 2) evaluating the expression pattern of *pCol-19::GFP* (referred to as “COL-19::GFP phenotype”; Fig. 2b).

**Figure 2.**
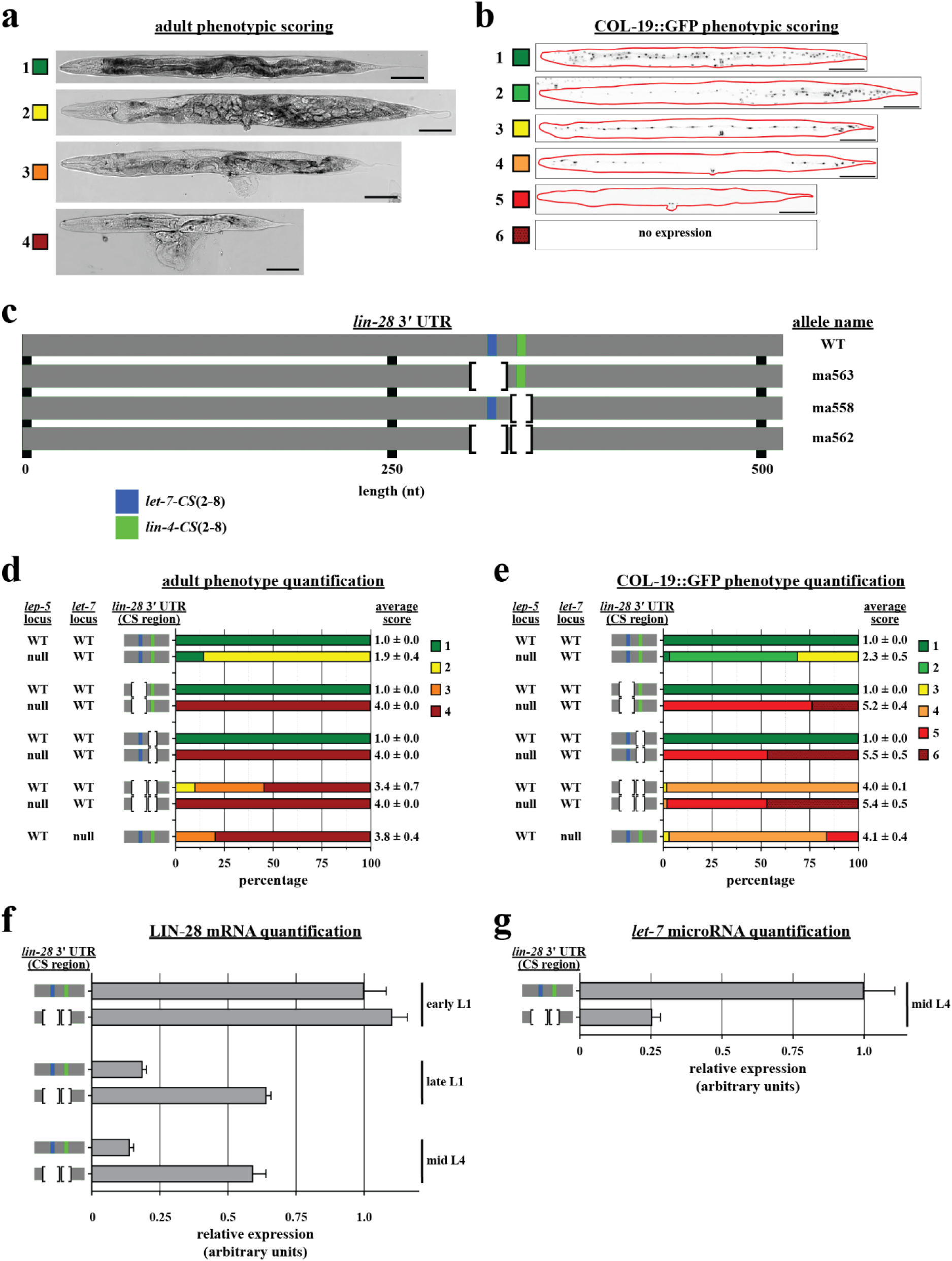
The loss of *let-7* and *lin-4* microRNA targeting of LIN-28 mRNA results in retarded development, which is exacerbated by the removal of the *lep-5* lncRNA. (a) Representative DIC images of retarded developmental phenotypes in adult *C. elegans* and the associated scores used for quantification. A score of 1 indicates adults that develop like wild-type, can actively lay eggs, and survive until a full brood is produced. A score of 2 indicates adults that are egg-laying defective and die due to a buildup of progeny within the mother. A score of 3 indicates an adult that dies due to bursting through a defective vulva after starting to produce progeny. A score of 4 indicates an adult that dies due to vulval bursting before producing progeny. Scale bars represent 100 µm. (b) Representative images of developmental phenotypes as seen in varying expression patterns of the adult-specific collagen reporter *pCol-19::GFP* and the score associated with each phenotype used for quantification. A score of 1 indicates a wild-type expression pattern in the hyp-7 syncytium, seam cells, and vulva cells. A score of 2 indicates partial expression in the hyp-7 syncytium while maintaining wild-type expression in seam and vulva cells. A score of 3 indicates little to no expression in hyp-7 with wild-type expression in seam and vulva cells. A score of 4 indicates little to no expression in hyp-7, partial expression in seam cells, and wild-type expression in vulva cells. A score of 5 indicates expression only in vulva cells. A score of 6 indicates no apparent expression in any cell. Scale bars represent 100 µm. (c) A depiction of the *C. elegans lin-28* 3’ UTR highlights the positions of the *let-7-CS* and the *lin-4-CS*. It presents the endogenous deletion alleles of the *let-7-CS* and the *lin-4-CS* used in the experimental procedures depicted in this manuscript. The allele *lin-28(ma563)* represents a deletion of the *let-7-CS*, the allele *lin-28(ma558)* denotes a deletion of the *lin-4-CS*, and the allele *lin-28(ma562)* encompasses deletions of both CSs. (d) Quantification of adult phenotypes was conducted in wild-type, *lin-28(ma563)*, *lin-28(ma558)*, *lin-28(ma562)*, and *let-7(ma393)* animals. All phenotypes were quantified under conditions with and without endogenous *lep-5*, except for *let-7(ma393)*, which was quantified only with *lep-5*. N’s (from top to bottom) = 44, 102, 58, 73, 60, 87, 79, 40, and 68. The data are presented as mean scores ± standard deviation. (e) Quantification of COL-19::GFP phenotypes was conducted in the same genotypes as described in panel (d). N’s (from top to bottom) = 42, 58, 72, 51, 88, 56, 46, 45, and 62. The data are presented as mean scores ± standard deviation. (f) RT-qPCR analysis of LIN-28 mRNA was conducted in whole animal extracts from early L1, late L1, and mid L4 stages of both wild-type and *lin-28(ma562)* genotypes. N = 4. Data are presented as mean ± standard deviation. (g) RT-qPCR analysis of *let-7* microRNA levels in whole animal extracts at the mid L4 stage for wild-type and *lin-28(ma562)* genotypes. N = 5. Data are presented as mean ± standard deviation.

To examine the effects of *let-7* and *lin-4* microRNA regulation on *C. elegans* LIN-28 expression, we employed CRISPR/Cas9 technology to delete the *let-7-CS(2-8)* and *lin-4-CS(2-8)* (hereafter referred to as the *let-7-CS* and *lin-4-CS*, unless otherwise specified) located within the *lin-28* 3’ UTR of the endogenous *lin-28* locus. We noted that the sequence flanking the *let-7-CS* exhibits high similarity to the *let-7-CS* sequence itself (CTACCA<u>CTACCTC</u>CTC; *let-7-CS* is underlined), suggesting that the deletion of sequences that are fully complementary to nucleotides 2-8 of the *let-7* microRNAs could potentially preserve a functional *let-7-CS*. Consequently, we also deleted the adjacent flanking sequence in conjunction with the CS to ensure complete ablation of *let-7* binding (*lin-28(ma563)*; hereafter referred to as *lin-28(Δlet-7-CS)*; Fig. 2c). In contrast, there was no redundant sequence flanking the *lin-4-CS*, and we replaced it—CTCAGGG—with TC (*lin-28(ma558)*; hereafter referred to as *lin-28(Δlin-4-CS)*), thereby introducing an EcoRI restriction enzyme site (Fig. 2c).

The single site deletions of either the *let-7-CS* or the *lin-4-CS* did not result in any observable phenotypes (Fig. 2d and 2e). However, the concurrent deletion of both CSs (*lin-28(ma562)*; hereafter referred to as *lin-28(ΔCSs)*) led to retarded developmental phenotypes, indicating that the *let-7-CS* and *lin-4-CS* function redundantly under standard laboratory conditions to negatively regulate LIN-28 expression (Fig. 2d and 2e).

MicroRNAs have the capacity to regulate the production of protein from an mRNA by inhibiting translation and/or by destabilizing the mRNA molecule (Shang et al. 2023). To investigate whether the targeting of *lin-28* by the *let-7* and *lin-4* microRNAs affects mRNA levels, we quantified LIN-28 mRNA using reverse transcription quantitative polymerase chain reaction (RT-qPCR) in wild-type and *lin-28(ΔCSs)* mutants during the early L1-stage (prior to the expression of either the *lin-4* or *let-7* microRNAs), the late L1-stage (following the increase in *lin-4* microRNAs levels), and the mid L4-stage (subsequent to the increases in the *let-7* microRNAs levels). We observed no significant differences in LIN-28 mRNA levels in early L1 samples of *lin-28(ΔCSs)* mutants relative to wild-type (Fig. 2f). However, an approximate three-fold increase in LIN-28 mRNA levels was detected in *lin-28(ΔCSs)* animals compared to wild-type at both the late L1 and mid L4 stages (Fig. 2f). These findings suggest that the *lin-4* and *let-7* microRNAs contribute to the destabilization of LIN-28 mRNA during the late L1 and subsequent stages.

In *C. elegans*, the *let-7* microRNA exhibits peak expression during the L4 developmental stage (Reinhart et al. 2000). The expression of the *let-7* microRNA at earlier developmental stages is inhibited by LIN-28; consequently, *lin-28* null (*lin-28(0)*) mutants undergo precocious development, which can be partially attributed to abnormally increased levels of the *let-7* microRNA beginning in the L2 stage (Ambros and Horvitz 1984; Johnson et al. 2003; Vadla et al. 2012). Therefore, we hypothesized that the misexpression of LIN-28 during later larval stages in *lin-28(ΔCSs)* animals would result in reduced *let-7* microRNA abundance compared to wild type at the L4 stage. Employing RT-qPCR, we found that *let-7* microRNA levels were approximately four-fold lower in *lin-28(ΔCSs)* in mid L4-stage larvae compared to wild-type, indicating that the misexpression of LIN-28 in *lin-28(ΔCSs)* represses *let-7* microRNA expression and suggesting that reduced *let-7* microRNA is likely an underlying cause of the retarded phenotypes observed in this mutant (Fig. 2g).

### The *let-7* and *lin-4* microRNAs collaborate with the *lep-5* lncRNA to downregulateLIN-28

We hypothesized that if the *let-7* and *lin-4* microRNAs were the sole negative regulators of LIN-28 expression, the gain-of-function phenotypes observed in *lin-28(ΔCSs)* animals would closely resemble the loss-of-function phenotypes of *let-7* null (*let-7(0)*) animals. However, *lin-28(ΔCSs)* animals exhibited relatively milder developmental delays compared to *let-7(0)* animals (Fig. 2d and 2e). These observations imply that an additional negative regulatory mechanism may be contributing to LIN-28 repression in later larval stages. As previously described, the *lep-5* lncRNA promotes LIN-28 degradation and has been proposed to constitute a mechanism for the removal of residual LIN-28 that is translated before the expression of the *let-7* and *lin-4* microRNAs (Kiontke et al. 2019). We postulated that, in addition to the removal of previously synthesized LIN-28, the *lep-5* lncRNA may also actively promote the degradation of nascent LIN-28 protein translated at later larval stages, thereby functioning in conjunction with the *let-7* and *lin-4* microRNAs to maintain low LIN-28 expression.

To investigate whether the *lep-5* lncRNA operates in conjunction with the *let-7* and *lin-4* microRNAs to downregulate LIN-28, we employed CRISPR/Cas9 technology to excise the entire *lep-5* sequence from the *C. elegans* genome (*lep-5(ma613)*; hereafter referred to as *lep-5(0)* unless otherwise noted). Consistent with previous findings, organisms deficient in *lep-5* exhibited relatively mild developmental retardation (Kiontke et al. 2019). However, *lin-28(ΔCSs)*; *lep-5(0)* double mutants exhibited significantly more pronounced developmental retardation than either single mutant, displaying phenotypes that were slightly more severe than *let-7(0)* mutants (Fig. 2d and 2e). Moreover, both *lin-28(Δlet-7-CS)*; *lep-5(0)* and *lin-28(Δlin-4-CS)*; *lep-5(0)* double mutants exhibited similarly profound developmental retardation that mirrored the phenotypes of *lin-28(ΔCSs)*; *lep-5(0)* double mutants (Fig. 2d and 2e). This indicates that the removal of either CS sensitizes developing *C. elegans* to further perturbations in LIN-28 expression.

Similar to *let-7(0)* mutants, which suffer essentially complete lethality at the L4 molt (and thus require balancing with a wild-type copy of *let-7* to facilitate propagation), *lin-28(Δlet-7-CS)*; *lep-5(0)*, *lin-28(Δlin-4-CS)*; *lep-5(0)*, and *lin-28(ΔCSs)*; *lep-5(0)* double mutants exhibit analogous lethality (and likewise must be balanced with a transgenic copy of *lep-5* expressed on an extrachromosomal array). Given the propensity for extrachromosomal arrays to be inherited unstably, we were able to assess the phenotypes of non-array-containing animals for these experiments.

### The *let-7* and *lin-4* microRNAs, along with the *lep-5* lncRNA, serve as the principal negative regulators of LIN-28 expression in seam and vulval cells

Loss of *lin-28* results in the precocious expression of seam and vulva cell fates normally associated with later developmental stages (e.g., adult-specific) (Ambros and Horvitz 1984; Euling and Ambros 1996). In contrast, opposite phenotypes—retarded development of seam and vulva cell fates—are observed in *lin-28* gain-of-function mutants (*lin-28(ΔCSs)* and *lep-5(0)*) (Fig. 2d and 2e). This supports the model that the normal temporal progression from larval to adult cell fates in these lineages depends on the progressive down regulation of LIN-28 throughout larval development. To assess the contributions of the *lin-4* and *let-7* microRNAs, as well as the *lep-5* lncRNA, in the temporal expression dynamics of endogenous LIN-28 protein in seam and vulva cells of wild-type animals, we quantified LIN-28 levels by utilizing a previously published endogenously tagged *lin-28::GFP* (Ilbay et al. 2021).

We observed that endogenously tagged LIN-28::GFP expression in the seam cells of wild-type animals was most prominent during the L1 larval stage and significantly diminished by the L4 stage (Fig. 3a and 3c). Likewise, developing vulval cells in L4 animals exhibited minimal LIN-28::GFP expression (vulval cells are not produced until after the L1 stage, thus precluding direct comparison to L4 vulval cells; Fig. 3b and 3d).

**Figure 3.**
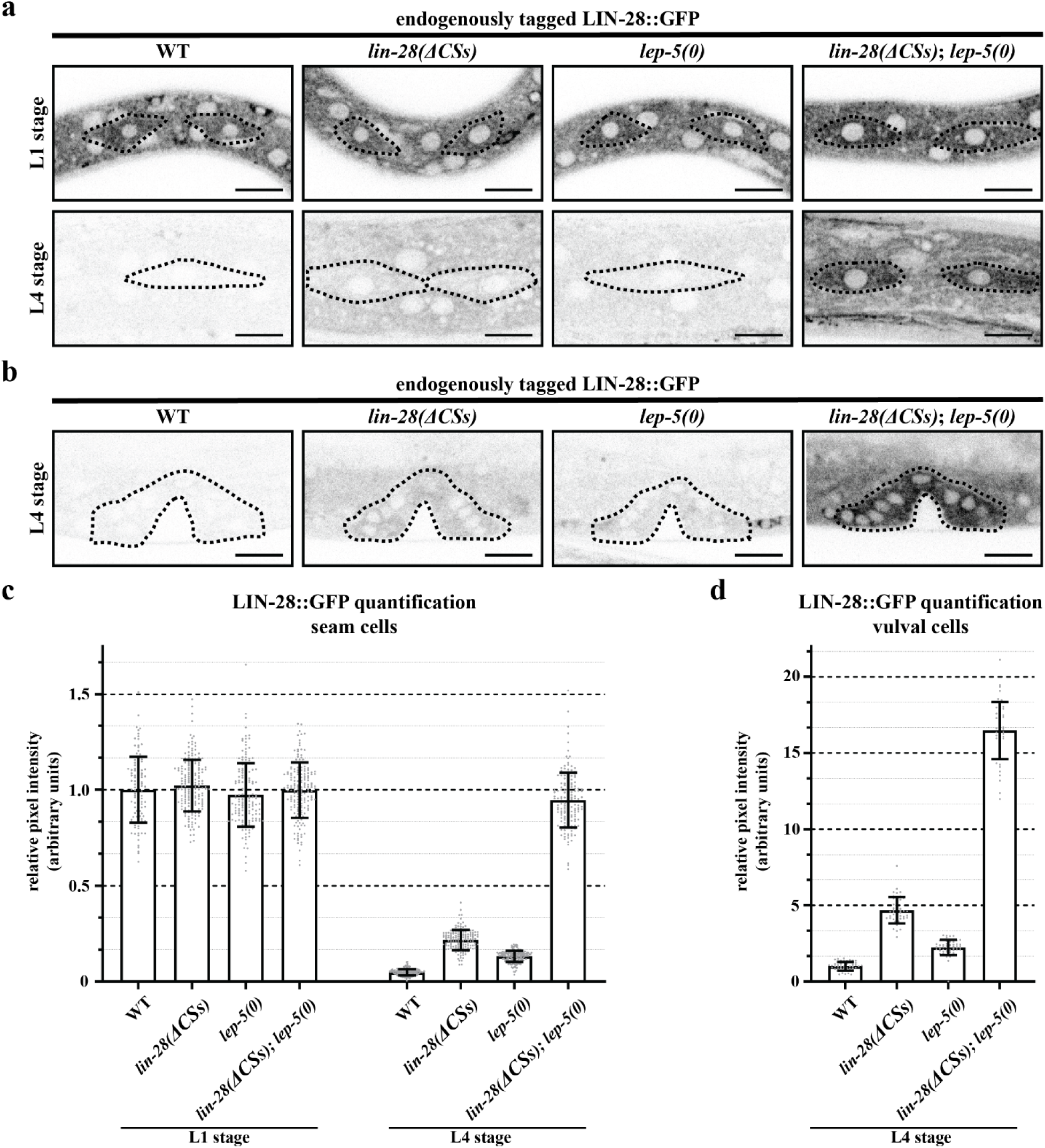
The *let-7* and *lin-4* microRNAs negatively regulate LIN-28 protein expression in a synergistic manner with *lep-5* lncRNA. (a) Representative images of endogenously tagged LIN-28::GFP in L1 and L4 wild-type, *lin-28(ΔCSs)*, *lep-5(0)*, and *lin-28(ΔCSs)*; *lep-5(0)* animals. Seam cells are outlined in black. Scale bars represent 10 µm. (b) Representative images of endogenously tagged LIN-28::GFP in L4 wild-type, *lin-28(ΔCSs)*, *lep-5(0)*, and *lin-28(ΔCSs)*; *lep-5(0)* animals. Vulva cells are outlined in black. Scale bars represent 10 µm. (c) Quantification of the average pixel intensity of LIN-28::GFP in seam cells described in panel (a). N’s (from left to right) = 108, 186, 156, 171, 147, 147, 165, and 153. Data are presented as mean ± standard deviation. (d) Quantification of the average pixel intensity of LIN-28::GFP in vulva cells described in panel (b). N’s (from left to right) = 46, 45, 54, and 47. Data are presented as mean ± standard deviation.

To assess LIN-28::GFP expression in animals deficient in the *let-7-CS* and *lin-4*-CS, we utilized CRISPR/Cas9 technology to generate the *lin-28(ΔCSs)* allele in *lin-28::GFP*-tagged animals, paralleling the modifications made in untagged animals. At the L1 stage, all the mutants examined (the *lin-28(ΔCSs)* and *lep-5(0)* single mutants, as well as the *lin-28(ΔCSs)*; *lep-5(0)* double mutants) exhibited nearly identical LIN-28::GFP expression levels in seam cells compared to wild-type (Fig. 3a and 3c). However, at the L4 stage, *lin-28(ΔCSs)* animals demonstrated an approximately 4.5-fold increase in LIN-28::GFP expression in seam cells and an approximately 4.7-fold increase in vulval cells relative to wild-type (Fig. 3c and 3d). *lep-5(0)* animals displayed an approximate 2.7-fold increase in seam cells and an approximate 2.2-fold increase in vulval cells at the L4 stage (Fig. 3a-3d). Strikingly, at the L4 stage, *lin-28(ΔCSs)*; *lep-5(0)* double mutant animals exhibited a much greater elevation in LIN-28::GFP levels, reflecting an approximate 19.4-fold de-repression in seam cells and an approximate 16.5-fold de-repression in vulva cells compared to their wild-type counterparts (Fig. 3a-3d).

These findings suggest a robust correlation between the penetrance of retarded developmental phenotypes and the levels of mis-expressed LIN-28. Furthermore, the dramatic synergy observed between the loss of microRNA and *lep-5* lncRNA regulation of LIN-28 (in terms of both developmental phenotypes and LIN-28::GFP mis-expression) indicates that the 3’ UTR-mediated microRNA regulation and the *lep-5* lncRNA mediated post-translational regulation of LIN-28 both exert major contributions to the developmental dynamics of LIN-28. Indeed, we note that the expression of LIN-28::GFP in *lin-28(ΔCSs)*; *lep-5(0)* double mutant animals at the L4 stage is similar to that observed at the L1 stage, underscoring the roles of the *let-7* and *lin-4* microRNAs, along with the *lep-5* lncRNA, as primary negative regulators of LIN-28 expression in seam and vulval cells.

### GFP tagging of LIN-28 mitigates the phenotypic manifestations associated with *lin-28* gain-of-function mutations

As described above, *lin-28(ΔCSs)* animals exhibit gain-of-function phenotypes that result in retarded development (Fig. 2d and 2e). Upon generating the *lin-28::GFP(ΔCSs)* strain, we observed that these animals did not display any apparent retarded developmental phenotypes (Fig. S1a). This observation suggests that the presence of the GFP tag on LIN-28 may alter the developmental dynamics of LIN-28 expression compared to untagged LIN-28, and/or that the tag may reduce the activity of the LIN-28 protein. To investigate the former possibility, we evaluated whether the GFP tag affected the L1-to-L4 downregulation of LIN-28 levels, by employing RT-qPCR to measure LIN-28 mRNA levels and western blotting with an anti-LIN-28 antibody to measure LIN-28 protein levels. The results indicated a subtle reduction in LIN-28::GFP mRNA during the early-L1 stage compared to untagged LIN-28 mRNA; however, no significant differences were observed in late-L1 and mid-L4 stages (Fig. S1b). Furthermore, we found no significant differences between untagged and GFP-tagged LIN-28 protein levels in L1 and L4 samples (Fig. S1c).

These findings indicate that the suppression of the gain-of-function phenotypes observed in *lin-28(ΔCSs)* animals with the presence of a GFP tag is attributed to a subtle hindrance of LIN-28 function (but not expression levels) imposed by the GFP tag. To assess the degree of reduction in LIN-28 activity conferred by the GFP tag, we examined *lin-28::GFP* larvae during larval stages for the phenotypic hallmarks associated with *lin-28* loss of function. All *lin-28::GFP* animals did not exhibit any discernible loss-of-function phenotypes and appeared indistinguishable from wild-type (Fig. S1a).

As previously mentioned, measuring *let-7* microRNA levels can serve as a sensitive molecular indicator of LIN-28 protein activity, given that *let-7* microRNA levels are negatively regulated by LIN-28. Accordingly, we measured *let-7* microRNA levels in L4 larvae using RT-qPCR and observed that *let-7* levels were approximately 1.6-fold greater in *lin-28::GFP* animals compared to their untagged counterparts, suggesting a slight reduction in LIN-28 function when tagged with GFP (Fig. S1d).

The above findings suggest that the GFP tag on LIN-28:GFP does not significantly impact LIN-28 developmental dynamics; however, it does result in a reduction of LIN-28 activity that is although subtle, nonetheless sufficient to suppress *lin-28* gain-of-function phenotypes. Notably, we observed a similar increase in LIN-28 mRNA levels when the CSs were removed from the 3’ UTR, irrespective of the presence of a GFP tag (Fig. S1e). Moreover, LIN-28:GFP activity is sufficient to cause full retarded phenotypes in the context of *lin-28::GFP(ΔCSs)*; *lep-5(0)* double mutants that were indistinguishable from those observed in their untagged counterparts (Fig. S1a). Furthermore, these mutants required balancing with an extrachromosomal array expressing transgenic *lep-5*, indicating that the removal of the primary negative regulators of LIN-28 expression (*let-7* microRNAs, *lin-4* microRNAs, and *lep-5* lncRNA) results in profound gain-of-function phenotypes, despite any suppression afforded by the GFP tag.

### The 3’ UTR of *lin-28* is non-essential under standard laboratory conditions and encompasses several positive regulatory elements

The *let-7-CS* and *lin-4-CS* sequences constitute 2.6% of the *C. elegans lin-28* 3’ UTR, indicating the distinct possibility for the existence of additional 3’ UTR regulatory elements. To investigate whether the *let-7-CS* and *lin-4-CS* are the predominant regulatory sequences in the *lin-28* 3’ UTR, we employed CRISPR/Cas9 technology to delete the entire *lin-28* 3’ UTR and replace it with a 64-bp segment of the *act-1* 3’ UTR, which retains the same polyadenylation signal (PAS) as the *lin-28* 3’ UTR (*lin-28(ma582)*; hereafter referred to as *lin-28(Δ3’ UTR)*; Fig. 4a). Unexpectedly, *lin-28(Δ3’ UTR)* animals exhibited a phenotypically wild-type appearance despite the absence of both the *let-7-CS* and *lin-4-CS* (Fig. 4b and 4c). This is in striking contrast to *lin-28(ΔCSs)* animals, where most of the 3’ UTR remains intact and pronounced *lin-28* gain-of-function phenotypes are observed. This finding suggests that the *lin-28* 3’ UTR contains sequences outside of the negative regulatory *let-7-CS* and *lin-4-CS* that appear to exert a largely balancing positive regulatory impetus.

**Figure 4.**
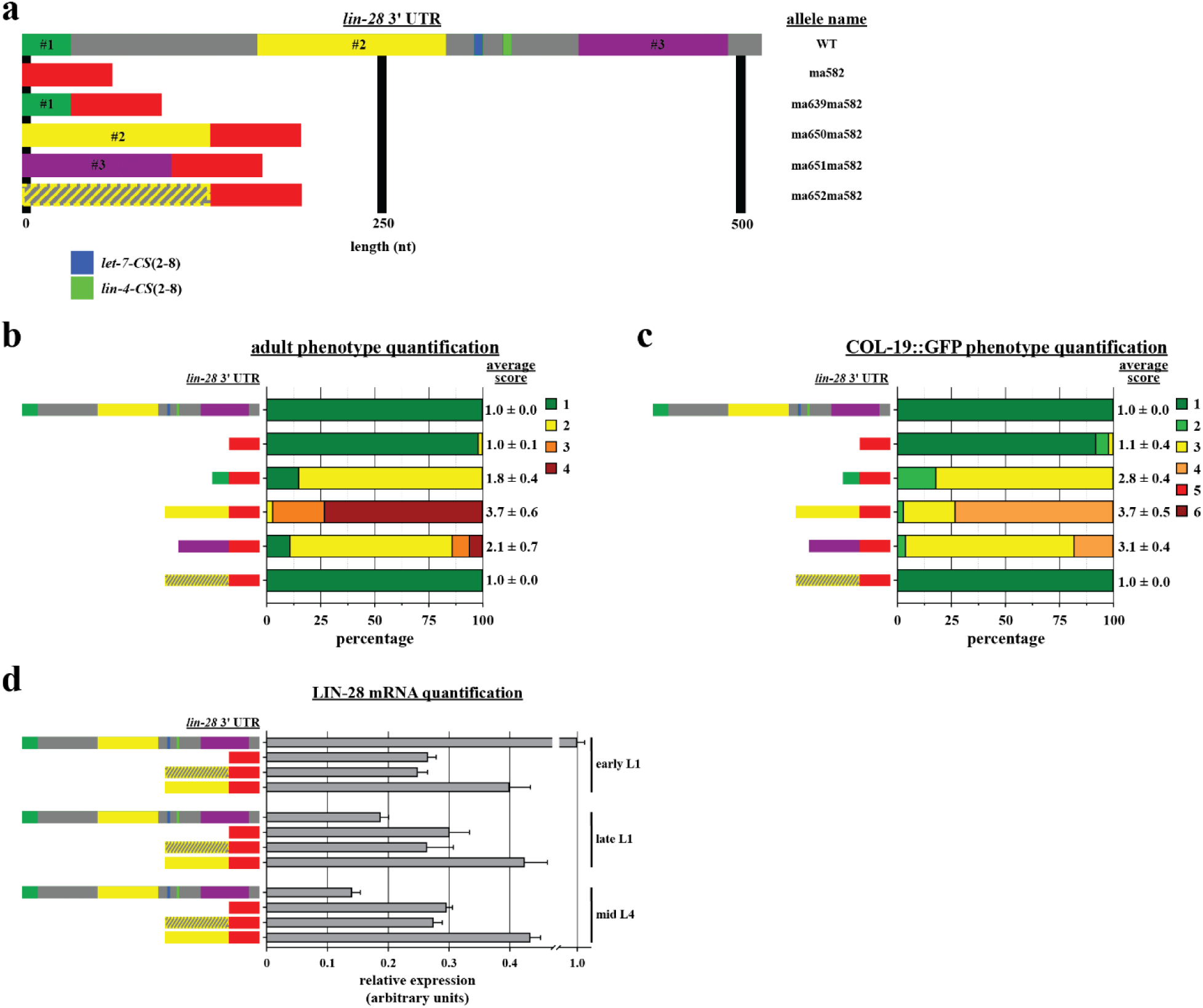
The *lin-28* 3’ UTR contains multiple positive regulatory sequences. (a) Depiction of the wild-type *C. elegans lin-28* 3’ UTR and the endogenous 3’ UTR mutant alleles used in experiments for this figure. *lin-28(ma582)* replaces the endogenous *lin-28* 3’ UTR with 64 bp of the *act-1* 3’ UTR. *lin-28(ma639ma582)* adds the piece highlighted in green from the endogenous *lin-28* 3’ UTR to *ma582*. *lin-28(ma650ma582)* adds the piece highlighted in yellow to *ma582*. *lin-28(ma651ma582)* adds the piece highlighted in magenta to *ma582*, and *lin-28(ma652ma582)* adds a scrambled version of *ma650* to *ma582*. (b) Quantification of adult phenotypes observed in the animals described in panel (a). Wild-type data are the same as shown in Fig. 2d. N’s (from top to bottom) = 44, 54, 125, 55, 96, and 72. Data are presented as mean score ± standard deviation. (c) Quantification of COL-19::GFP phenotypes observed in the same genotypes as panel (b). Wild-type data are the same as shown in Fig. 2e. N’s (from top to bottom) = 42, 51, 71, 55, 51, and 64. Data are presented as mean score ± standard deviation. (d) RT-qPCR analysis of LIN-28 mRNA in whole animal extracts from early L1, late L1, and mid L4 stages of wild-type, *lin-28(ma582)*, *lin-28(ma652ma582)*, and *lin-28(ma650ma582)* genotypes. Wild-type data are the same as shown in Fig. 2f. N = 4. Data are presented as mean ± standard deviation.

To investigate whether these positive regulatory elements contribute to the misexpression of LIN-28 in the absence of the *let-7-CS* and *lin-4-CS*, we employed CRISPR/Cas9 technology to generate a series of strains with deletions that systematically eliminated portions of the endogenous *lin-28* 3’ UTR. We then assessed these strains for retarded developmental phenotypes (Fig. S2a and S2b). Our analysis identified three specific regions within the *lin-28* 3’ UTR whose presence correlated with the expression of retarded phenotypes in constructs lacking both the *let-7-CS* and *lin-4-CS*, indicating that these regions likely possess positive regulatory functions (Fig. S2b). Notably, all 3’ UTR truncation mutants that retained the CSs exhibited wild-type phenotypes, suggesting that *let-7* and *lin-4*-mediated repression of LIN-28 does not require an intact 3’ UTR and is sufficient to counteract any promotion of LIN-28 expression conferred by positive elements within the 3’ UTR (Fig. S2b).

To test whether the three positive regulatory regions identified from our *lin-28* 3’ UTR truncation experiments were adequate to promote LIN-28 misexpression in the absence of any residual endogenous *lin-28* 3’ UTR sequences, we employed CRISPR/Cas9 technology to insert each respective region at the beginning of the artificial 3’ UTR of *lin-28(Δ3’ UTR)* animals and assessed them for developmental phenotypes indicative of LIN-28 misexpression (*lin-28(ma639ma582)*, *lin-28(ma650ma582)*, *lin-28(ma651ma582)*, and *lin-28(ma652ma582)*; Fig. 4a). The inclusion of each individual region resulted in varying degrees of developmental retardation when compared to a scrambled control, which exhibited characteristics indistinguishable from wild-type (Fig. 4b and 4c). These findings indicate that each region is sufficient to promote LIN-28 expression in the absence of the rest of the *lin-28* 3’ UTR.

As previously indicated, *lin-28(ΔCSs)* animals exhibit elevated levels of LIN-28 mRNA at later stages in comparison to wild-type animals, likely due to a reduction in mRNA turnover mediated by the *let-7* and *lin-4* microRNAs (Fig. 2f). Although *lin-28(Δ3’ UTR)* animals display a wild-type phenotype, they also lack the *let-7-CS* and *lin-4*-*CS* and, therefore, might be expected to exhibit milder downregulation of LIN-28 mRNA compared to wild type. To investigate the dynamics of LIN-28 mRNA in *lin-28(Δ3’ UTR)* animals, we employed RT-qPCR to quantify LIN-28 mRNA levels in *lin-28(Δ3’ UTR)* mutants at early L1, late L1, and mid L4 larval stages, and observed approximately the same levels of LIN-28 mRNA at all time points, which is consistent with 3’ UTR-mediated down regulation of LIN-28 mRNA (Fig. 4d). However, the levels of LIN-28 mRNA at all stages in *lin-28(Δ3’ UTR)* mutants were approximately 25% of those observed in wild-type and *lin-28(ΔCSs)* animals at the early L1 stage, and approximately half of the levels seen in *lin-28(ΔCSs)* animals at the late L1 and mid L4 stages (Fig. 1F and 4d). This latter observation indicates that *lin-28* 3’ UTR sequences outside of the *let-7-CS* and *lin-4*-*CS* promote LIN-28 mRNA accumulation at all stages.

The observation that removal of the *lin-28* 3’ UTR resulted in decreased LIN-28 mRNA levels compared to *lin-28(ΔCSs)* animals prompted us to hypothesize that the positive elements enhance LIN-28 expression through the stabilization of its mRNA. To investigate this hypothesis, we utilized RT-qPCR to measure LIN-28 mRNA levels at early L1, late L1, and mid L4 larval stages in the strain in which the most potent positive element was inserted into the 3’ UTR of *lin-28(Δ3’ UTR)* animals (*lin-28(ma650ma582)*; hereafter referred to as *lin-28(positive #2::Δ3’ UTR)*; Fig. 4a). In these *lin-28(positive #2::Δ3’ UTR)* animals, LIN-28 mRNA levels remained approximately constant across all three timepoints, representing about 40% of wild-type early L1 levels, and approximately twice that of wild-type levels at late L1 and mid L4 timepoints (Fig. 4d). Furthermore, at all three timepoints, LIN-28 mRNA levels were approximately one-third higher than both *lin-28(Δ3’ UTR)* and *lin-28(scrambled::Δ3’ UTR)* mutants (Fig. 4d). These results support the conclusion that the positive elements within the *lin-28* 3’ UTR facilitate LIN-28 expression by stabilizing its mRNA.

### The 3’ UTR mediated regulation of *lin-28* functions independently of *lep-5*-mediated post-translational regulation

The essentially wild-type phenotype observed in *lin-28(Δ3’ UTR)* animals suggests that removal of the entire endogenous *lin-28* 3’ UTR releases LIN-28 expression from 3’ UTR-mediated regulation by balancing negative regulation (mediated by the *lin-4-CS* and *let-7*-*CS*) and positive regulation (mediated by the aforementioned positive elements). This observation raises the question of what mechanism(s) is/are responsible for LIN-28 downregulation in the absence of these balancing regulatory influences associated with the 3’ UTR, thereby facilitating the nearly wild-type development of *lin-28(Δ3’ UTR)* animals. We noticed that *lin-28(Δ3’ UTR)* animals displayed very mild developmental retardation, suggesting that the combined deletion of negative and positive regulatory elements leads to a slight misexpression of LIN-28 in these animals. To test whether the slight misexpression LIN-28 in *lin-28(Δ3’ UTR)* animals could be aggravated by the removal of the *lep-5* lncRNA, we assessed the developmental phenotypes in *lin-28(Δ3’ UTR)*; *lep-5(0)* double mutants. These double mutants exhibited severe developmental retardation that was indistinguishable from that observed in *lin-28(ΔCSs)*; *lep-5(0)* double mutants (Fig. 5b and 5c).

**Figure 5.**
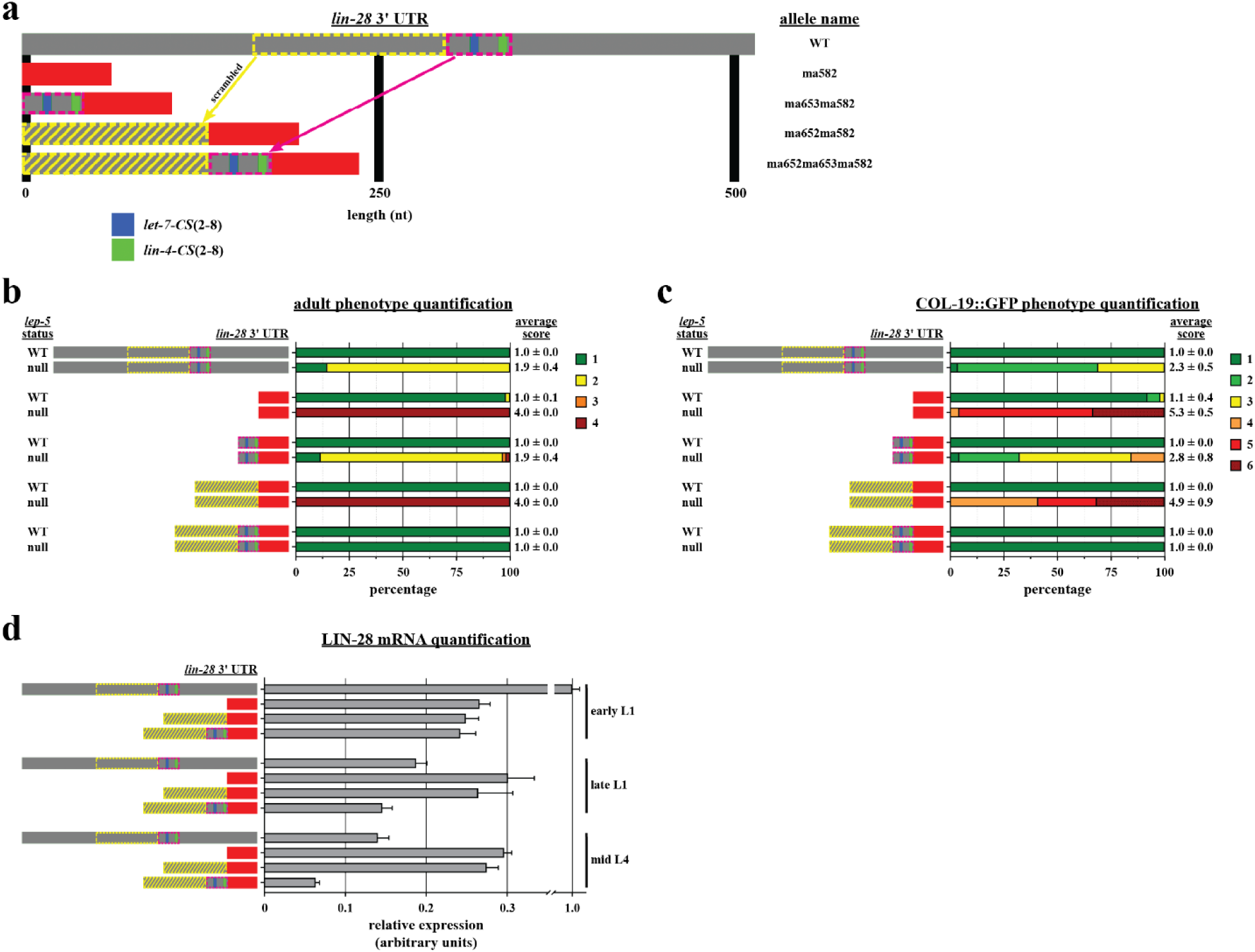
The *lin-28* 3’ UTR is non-essential under standard laboratory conditions. (a) Depiction of the wild-type *C. elegans lin-28* 3’ UTR and the endogenous 3’ UTR mutant alleles used in experiments for this figure. *lin-28(ma582)* replaces the endogenous *lin-28* 3’ UTR with 64 bp of the *act-1* 3’ UTR. *lin-28(ma653ma582)* adds a segment of the endogenous *lin-28* 3’ UTR containing the *let-7-CS* and *lin-4-CS* to *ma582*. *lin-28(ma652ma582)* adds a scrambled spacer before *ma582*, and *ma652ma653ma582* adds both the scrambled spacer and CSs to *ma582*. (b) Quantification of adult phenotypes observed in the animals described in panel (a). All phenotypes were quantified in the presence and absence of endogenous *lep-5*. Wild-type 3’ UTR data are the same as shown in Fig. 2d. *lin-28(ma582)* and *lin-28(ma652ma582)* data in the presence of wild-type *lep-5* are the same as shown in Fig. 4b. N’s (from top to bottom) = 44, 102, 54, 43, 142, 120, 72, 62, 98, and 142. Data are presented as mean ± standard deviation. (c) Quantification of COL-19::GFP phenotypes observed in the same genotypes as panel (b). Wild-type 3’ UTR data are the same as shown in Fig. 2e. *lin-28(ma582)* and *lin-28(ma652ma582)* data in the presence of wild-type *lep-5* are the same as shown in Fig. 4c. N’s (from top to bottom) = 42, 58, 51, 48, 60, 71, 64, 95, 84, and 82. Data are presented as mean ± standard deviation. (d) RT-qPCR analysis of LIN-28 mRNA in whole animal extracts from early L1, late L1, and mid L4 stages of wild-type, *lin-28(ma582)*, *lin-28(ma652ma582)*, and *lin-28(ma652ma653ma582)* genotypes. Wild-type 3’ UTR data are the same as shown in Fig. 2f. N = 4. Data are presented as mean ± standard deviation.

Similar to previously described double mutants lacking *lep-5*, the maintenance of the *lin-28(Δ3’ UTR)*; *lep-5(0)* double mutant genotype necessitated the use of a transgene expressing *lep-5* on an extrachromosomal array. This significant enhancement of *lin-28(Δ3’ UTR)* phenotypes by loss of *lep-5* suggests that the normal developmental dynamics of LIN-28 reflect contributions from 3’ UTR-mediated regulation of LIN-28 synthesis, and 3’ UTR-independent regulation of LIN-28 stability by the *lep-5* lncRNA.

### The *let-7-CS* and *lin-4-CS* can function independently of the rest of the *lin-28* 3’ UTR

The severe *lin-28* misexpression phenotypes of *lin-28(Δ3’ UTR)*; *lep-5(0)* double mutants provided an opportunity to experimentally test whether the *let-7-CS* and the *lin-4-CS* can function to downregulate LIN-28 in the absence of other 3’ UTR sequences. Accordingly, we employed CRISPR/Cas9 to insert a 42-bp sequence derived from the wild-type *lin-28* 3’ UTR, encompassing both the *let-7-CS* and the *lin-4-CS*, at the beginning of the artificial 3’ UTR in *lin-28(Δ3’ UTR)* animals (*lin-28(ma653ma582)*; hereafter referred to as *lin-28(CSs::Δ3’ UTR)*; Fig. 5a) and investigated whether this 42 bp sequence was sufficient to suppress the severely retarded developmental phenotypes observed in *lin-28(Δ3’ UTR)*; *lep-5(0)* double mutant animals. The *lin-28(CSs::Δ3’ UTR)*; *lep-5(0)* animals demonstrated a significant but incomplete suppression of the gain-of-function phenotypes observed in *lin-28(Δ3’ UTR)*; *lep-5(0)* animals (Fig. 5b and 5c). This result indicates that the *lin-28* 3’ UTR CSs can downregulate LIN-28 independently of other 3’ UTR sequences, suggesting that the mechanism of microRNA-mediated repression of LIN-28 does not necessarily involve overcoming the activity of the positive regulatory elements

The incomplete suppression of the *lin-28(Δ3’ UTR)*; *lep-5(0)* phenotypes in *lin-28(CSs::Δ3’ UTR)*; *lep-5(0)* animals led us to hypothesize that the 42 bp *let-7-CS* and the *lin-4-CS* element may not be positioned optimally for full microRNA activity, given their proximity to the stop codon of the *lin-28* gene. Therefore, we designed a 130-bp spacer consisting of a scrambled segment of the endogenous *lin-28* 3’ UTR and employed CRISPR/Cas9 technology to insert it between the *lin-28* stop codon and the CSs of *lin-28(CSs::Δ3’ UTR)* animals (*lin-28(ma652ma653ma582)*; henceforth referred to as *lin-28(spacer::CSs::Δ3’ UTR)*) and a control line (*lin-28(ma652ma582)*; hereafter referred to as *lin-28(spacer::Δ3’ UTR)*; Fig. 5a). The insertion of the spacer in the absence of the CSs did not result in significant suppression of the gain-of-function phenotypes of *lin-28(Δ3’ UTR)*; *lep-5(0)* (Fig. 5b and 5c). However, the inclusion of the spacer upstream of the CSs resulted in complete suppression, thereby reinforcing the conclusion that the *let-7-CS* and *lin-4-CS* can function independently of the rest of the native *lin-28* 3’ UTR sequences (Fig. 5b and 5c). Additionally, this suggests that positioning of the CSs further from the *lin-28* stop codon is more optimal for *let-7* and *lin-4-*mediated repression of LIN-28 expression.

In the absence of other *lin-28* 3’ UTR sequences, particularly the positive elements, one might have anticipated that the net effect of restoring *let-7* and *lin-4* microRNA repression in the *lin-28(spacer::CSs::Δ3’ UTR)* construct could result in precocious phenotypes characteristic of *lin-28* loss-of-function. However, *lin-28(spacer::CSs::Δ3’ UTR)* animals did not exhibit any discernible loss-of-function phenotypes and appeared phenotypically indistinguishable from wild-type (Fig. 5b and 5c).

Based on our previous findings showing that the deletion of the *let-7* and *lin-4* CSs from the *lin-28* 3’ UTR stabilizes LIN-28 mRNA (Fig. 2f), we hypothesized that the addition of these CSs in *lin-28(spacer::CSs::Δ3’ UTR)* animals should destabilize LIN-28 mRNA when the microRNAs are expressed. Using RT-qPCR, we observed decreased levels of LIN-28 mRNA in late L1 and mid L4 samples compared to both wild-type and controls lacking the CSs (Fig. 5d). This suggests that the *let-7* and *lin-4* microRNAs are sufficient to repress LIN-28 expression by destabilizing LIN-28 mRNA independently of rest of the *lin-28* 3’ UTR.

### The deletion of the entire *lin-28* 3’ UTR leads to relatively mild, yet significant, alterations in the developmental dynamics of LIN-28 expression

Based on our RT-qPCR data indicating a reduction of LIN-28 mRNA at the L1 stage and an elevation at the L4 stage in *lin-28(Δ3’ UTR)* larvae compared to wild-type, we hypothesized that LIN-28 protein levels would be likewise decreased in *lin-28(Δ3’ UTR)* L1 larvae and elevated in L4 larvae. To test this hypothesis, we utilized CRISPR/Cas9 technology to generate *lin-28::GFP(Δ3’ UTR)* animals. As predicted, LIN-28::GFP expression was approximately 50% lower in the seam cells of L1 larvae and approximately threefold higher in both the seam and vulval cells of L4 larvae in *lin-28::GFP(Δ3’ UTR)* animals when compared to wild-type (Fig. 6a-6d). These results provide support for the model that the *lin-28* 3’ UTR mediates positive and negative regulatory inputs resulting in the promotion of LIN-28 expression during early developmental stages, and the repression of LIN-28 expression in later stages.

**Figure 6.**
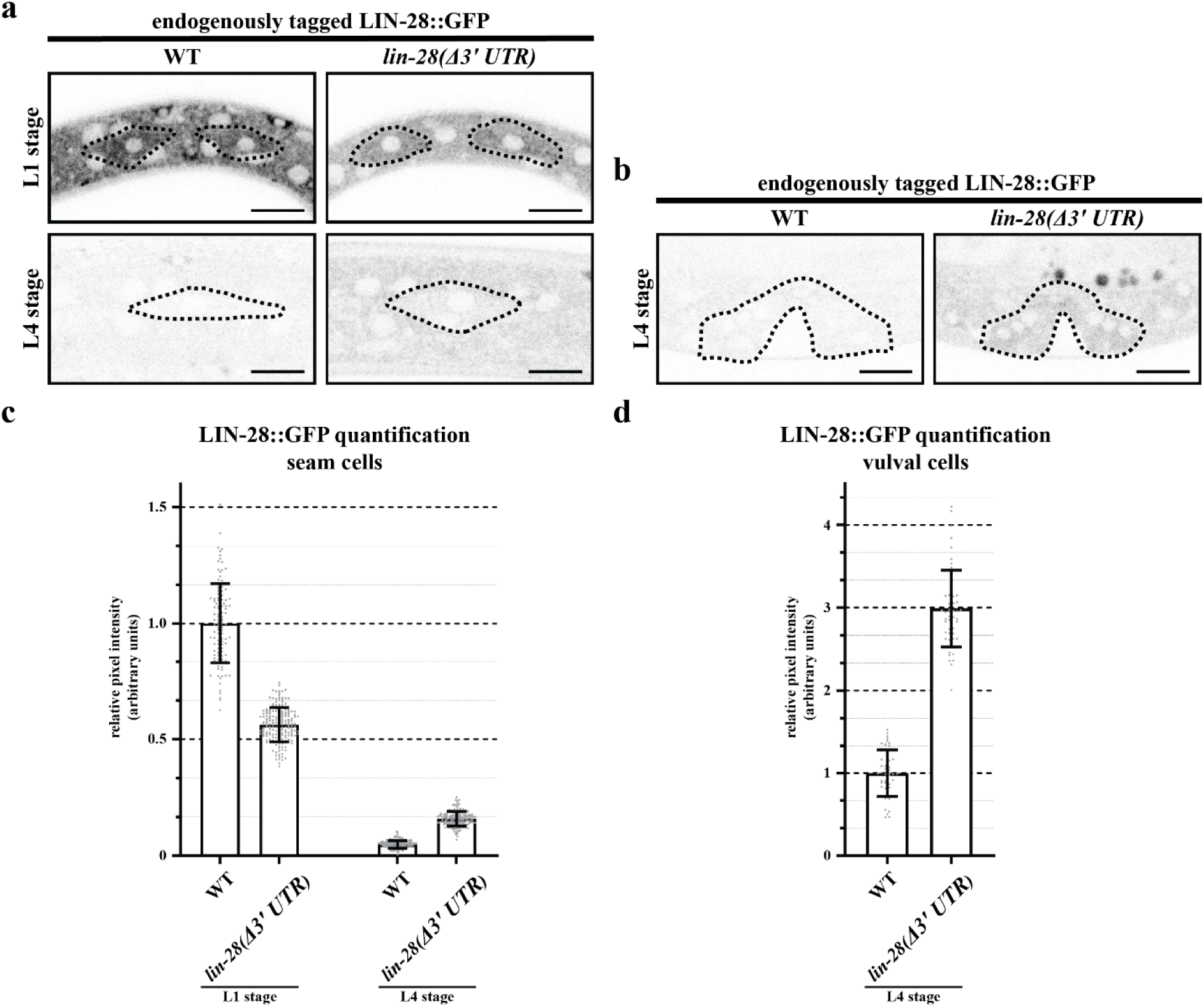
The 3’ UTR of *lin-28* enhances the expression of LIN-28::GFP during early development. (a) Representative images of endogenously tagged LIN-28::GFP in L1 and L4 wild-type and *lin-28(Δ3’ UTR)* animals. Wild-type images are the same as shown in Fig. 3a. Seam cells are outlined in black. Scale bars represent 10 µm. (b) Representative images of endogenously tagged LIN-28::GFP in L4 wild-type and *lin-28(Δ3’ UTR)* animals. Wild-type images are the same as shown in Fig. 3b. Vulva cells are outlined in black. Scale bars represent 10 µm. (c) Quantification of the average pixel intensity of LIN-28::GFP in seam cells described in panel (a). Wild-type data are the same as shown in Fig. 3c. N’s (from left to right) = 108, 183, 165, and 165. Data are presented as mean ± standard deviation. (d) Quantification of the average pixel intensity of LIN-28::GFP in vulva cells described in panel (b). Wild-type data are the same as shown in Fig. 3d. N’s (from left to right) = 46 and 52. Data are presented as mean ± standard deviation.

### The heterochronic phenotypes of *lin-28* 3’ UTR mutants exhibit sensitivity to development at specific temperatures or in the presence of the pathogen *Pseudomonas aeruginosa*

In *C. elegans*, *let-7* family microRNAs are expressed from the *mir-48*, *mir-84*, *mir-241*, and *let-7* loci during progression from the L2-through adult stages of development. miR-48, miR-84, and miR-241 are the dominant family members expressed in the L2 and L3 stages, while the *let-7* microRNA is primarily up regulated in the L4 stage. Therefore, much of the *let-7* family-mediated down regulation of LIN-28, which occurs by the L2 stage, is exerted by the combined action of miR-48, miR-84, miR-241. Accordingly, loss-of-function mutants of *mir-48*, *mir-84*, and *mir-241* exhibit retarded heterochronic phenotypes that are analogous to those observed for LIN-28 misexpression mutants examined in this study (Abbott et al. 2005).

Remarkably, the retarded developmental phenotypes of *mir-48*, *mir-84*, and *mir-241* compound mutants can be modified by environmental conditions. For example, exposure of *mir-48*, *mir-84*, and *mir-241* mutants to the pathogenic bacterium *Pseudomonas aeruginosa* exacerbates their retarded phenotypes, demonstrating that in animals lacking these microRNAs, gene regulatory mechanisms specifying temporal cell fates are sensitive to environmental stresses associated with bacterial infection (Ren and Ambros 2015).

To test if the heterochronic phenotypes of *lin-28* 3’ UTR mutants would likewise exhibit sensitivity to exposure to *P. aeruginosa*, we cultured wild-type, *lin-28(ΔCSs)*, and *lin-28(Δ3’ UTR)* larvae on the pathogenic *P. aeruginosa* strain PA14 and the non-pathogenic GacA mutant of PA14. *C. elegans* larvae and young adults are tolerant of PA14 as a food source; however, exposure to PA14 is lethal to older adults, which precluded the reliable assessment of adult morphological phenotypes (Tan et al. 1999; Ren and Ambros 2015; Mirza et al. 2023). Nevertheless, we were able to assess *pCol-19::GFP* expression by scoring young adults. Wild-type animals exhibited no significant differences in *pCol-19::GFP* expression when exposed to either PA14 or GacA (Fig. 7a). In contrast, the retarded developmental *pCol-19::GFP* phenotypes observed in *lin-28(ΔCSs)* animals were significantly more severe upon exposure to PA14 compared to GacA (Fig. 7a). Notably, *lin-28(Δ3’ UTR)* animals, which displayed wild-type-like *pCol-19::GFP* expression when exposed to GacA, exhibited retarded *pCol-19::GFP* expression patterns when exposed to PA14 (Fig. 7a). This finding indicates that the loss of regulation of LIN-28 by the *let-7* and *lin-4* microRNAs sensitizes larval cell fate progression to stressors associated with exposure to virulent *P. aeruginosa*.

**Figure 7.**
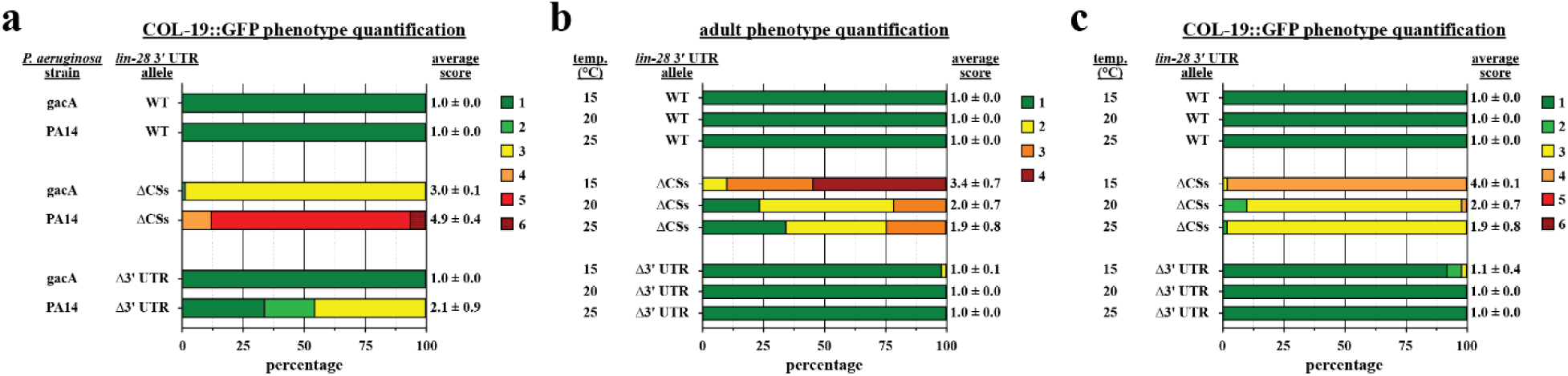
*lin-28* 3’ UTR mutants exhibit increased sensitivity to environmental stressors. (a) Quantification of COL-19::GFP phenotypes of wild-type, *lin-28(Δ3’ UTR)*, and *lin-28(ΔCSs)* strains exposed to the non-pathogenic *P. aeruginosa* strain GacA and the pathogenic *P. aeruginosa* strain PA14. N’s (from top to bottom) = 98, 44, 66, 49, 107, and 44. Data are presented as mean score ± standard deviation. (b) Quantification of adult phenotypes of wild-type, *lin-28(Δ3’ UTR)*, and *lin-28(ΔCSs)* strains grown at 15°C, 20°C, and 25°C. 15°C wild-type and *lin-28(ΔCSs)* data are the same as shown in Fig. 2d. 15°C *lin-28(Δ3’ UTR)* data are the same as shown in Fig. 4b. N’s (from top to bottom) = 44, 63, 41, 79, 140, 90, 54, 69, and 62. Data are presented as mean score ± standard deviation. (c) Quantification of COL-19::GFP phenotypes observed in the same genotypes as panel (b). 15°C wild-type and *lin-28(ΔCSs)* data are the same as shown in Fig. 2e. 15°C *lin-28(Δ3’ UTR)* data are the same as shown in Fig. 4c. N’s (from top to bottom) = 42, 63, 53, 46, 50, 52, 51, 47, and 62. Data are presented as mean score ± standard deviation.

All previous phenotypic experiments described here that quantified the penetrance of retarded developmental phenotypes were conducted at 15°C. However, the SK conditions necessary for exposure to PA14 had to be performed at 25°C. During these experiments, we observed that the developmental phenotypes of *lin-28(ΔCSs)* animals raised on GacA at 25°C were not as severe compared to previous experiments in which animals were raised on *E. coli* at 15°C. To assess whether *lin-28(ΔCSs)* and *lin-28(Δ3’ UTR)* animals exhibited sensitivity to temperature, we raised each strain on *E. coli* at 15°C, 20°C, and 25°C, and compared their phenotypes to the wild-type. No significant differences in phenotypes across the tested temperatures were observed for wild-type and *lin-28(Δ3’ UTR)* animals (Fig. 7b and 7c). In contrast, *lin-28(ΔCSs)* animals exhibited more penetrant retarded developmental phenotypes when raised at 15°C compared to those raised at 20°C and 25°C (Fig. 7b and 7c).

## DISCUSSION

In this study, we demonstrate that across all currently characterized *Caenorhabditis* species, *lin-4* microRNA-mediated repression of LIN-28 is conserved, while *let-7* microRNA-mediated repression is predominantly restricted to the *Elegans* and *Japonica* groups of *Caenorhabditis* species. We established that a *let-7-CS* and a *lin-4-CS* located within the *C. elegans lin-28* 3’ UTR function in a semi-redundant manner to repress LIN-28 by regulating both mRNA abundance and inhibiting translation. Furthermore, these elements act synergistically with the post-translational repression of LIN-28 mediated by the *lep-5* lncRNA. Our findings also revealed that deletion of the entire *lin-28* 3’ UTR results in animals exhibiting a near wild-type phenotype, indicating that the *lin-28* 3’ UTR contains cis-acting elements that facilitate the positive regulation of LIN-28 expression and counteract the repressive effects of the *let-7* and *lin-4* microRNAs. Our data show that these positive acting elements function by stabilizing LIN-28 mRNA. Finally, we determined that despite their wild-type appearance, *lin-28* 3’ UTR mutants display sensitivity to environmental conditions: development of *lin-28* 3’ UTR mutants at alternative culture temperatures or in the presence of the pathogenic bacterium *P. aeruginosa* elicited cryptic heterochronic phenotypes. We conclude that the multiple layers of stringent regulation of LIN-28 expression described here— encompassing mRNA stability, protein synthesis, and protein stability combine to ensure that LIN-28 is accurately and precisely downregulated at the appropriate time, irrespective of environmental conditions.

In *C. elegans* and in mammals, LIN-28 functions as a negative regulator of the *let-7* microRNA, such that the loss of LIN-28 results in elevated levels of the *let-7* microRNA (Johnson et al. 2003; Viswanathan et al. 2008; Heo et al. 2008). Furthermore, the 3’ UTR of LIN-28 mRNA in both *C. elegans* and mammals contain *let-7-CSs*, indicating a highly conserved mutual negative regulatory relationship between *let-7* microRNAs and LIN-28 (Reinhart et al. 2000; Moss and Tang 2003). In many of the *Caenorhabditis* species examined in this study, *lin-28* is predicted to be directly regulated by *let-7* family microRNAs, as well as *lin-4* microRNA. Interestingly, in some of the *Caenorhabditis* species, *lin-28* is predicted to be regulated by the *lin-4* microRNA but not by the *let-7* microRNAs.

Our analysis of *lin-28* 3’ UTR sequence conservation among *Caenorhabditis* species revealed two types of *lin-4-CSs*—specifically, 2-8 pairing and 3-8 pairing. Notably, nucleotides 2-4 of the *lin-4* microRNA seed sequence consist of a triplet of guanines, allowing for three potential seed pairings with a guanine that could be bulged at positions 2, 3, or 4 (for clarity, we will continue to refer to these sites as *lin-4-CS(3-8)*). Our analysis indicated that the number of *lin-4-CS(3-8)* sites was significantly greater than that of *lin-4-CS(2-8)* sites among all analyzed *Caenorhabditis lin-28* 3’ UTRs (65 sites versus 35 sites, respectively). The conservation of this alternative pairing suggests that the utilization of *lin-4-CS(3-8)* sites may confer distinct functional characteristics in comparison to *lin-4-CS(2-8)* sites.

Deletion of either the *let-7-CS* alone or the *lin-4-CS* alone from the *C. elegans lin-28* 3’ UTR resulted in animals that exhibited a wild-type phenotype under standard laboratory conditions. However, perturbation of LIN-28 expression by removing the post-translational negative regulator *lep-5* lncRNA in these single site deletion mutants led to highly penetrant retarded developmental phenotypes, indicative of extreme LIN-28 misexpression. These findings suggest that these microRNA sites function semi-redundantly, as the loss of either single site renders the animal susceptible to further alterations in LIN-28 expression.

It has been previously demonstrated that the *lep-5* lncRNA, which is upregulated during the L2 stage, elicits post-translational degradation of LIN-28 protein, including LIN-28 synthesized prior to the expression of the *lin-4* and *let-7* microRNAs following the L1 stage (Kiontke et al. 2019). If the only function of the *lep-5* lncRNA was to degrade LIN-28 translated during the L1 stage, and not during subsequent stages, one would expect an additive effect on LIN-28 levels when comparing the *lin-28(ΔCSs)* and *lep-5(0)* single and double mutants. However, our findings reveal a synergistically elevated expression of LIN-28 in double mutants, as evidenced by the severe phenotypes manifested in *lin-28(ΔCSs)*; *lep-5(0)* double mutants, compared to the relatively mild retarded developmental phenotypes observed in *lin-28(ΔCSs)* and *lep-5(0)* single mutants. This is further corroborated by the extent of LIN-28::GFP expression in single versus double mutants; in L4 animals, *lin-28(ΔCSs)* and *lep-5(0)* single mutants exhibit approximately fourfold and fivefold upregulation of LIN-28::GFP expression in seam cells, respectively, compared to approximately twentyfold upregulation in double mutants. These results suggest that the *lep-5* lncRNA functions continuously throughout the L2 to L4 stages as an attenuator of LIN-28 accumulation, thereby complementing microRNA-mediated inhibition of LIN-28 synthesis.

MicroRNAs are known for their role in modulating the level of protein synthesis from target mRNAs through the repression of target mRNA stability and/or the inhibition of target mRNA translational efficiency (TE) (Cottrell et al. 2017). In the context of the regulation of *lin-28* by the *let-7* and *lin-4* microRNAs, our findings do not conclusively elucidate the contributions of mRNA stability and TE regulation; however, they suggest the potential involvement of both mechanisms. Our quantification of LIN-28::GFP upregulation in seam cells at the L4 stage of *lep-5(0)* animals, with and without the CSs in the *lin-28* 3’ UTR, indicates an approximate seven-fold effect of the *let-7* and *lin-4* microRNAs on LIN-28 levels. At the whole animal level, LIN-28 mRNA expression is upregulated by approximately four-fold upon deletion of the CSs from the *lin-28* 3’ UTR. Since the *lep-5* lncRNA does not affect LIN-28 mRNA levels, this four-fold increase can be interpreted as being mediated by microRNAs (Kiontke et al. 2019). A limitation of these interpretations lies in the comparison of whole animal mRNA measurements to protein measurements in seam cells, and it is possible that regulatory mechanisms may differ among distinct cell types. Furthermore, the impact of microRNAs on LIN-28 mRNA levels may also be complicated by potential autoregulatory effects exerted by LIN-28 protein on LIN-28 mRNA (Stefani et al. 2015).

Previous studies investigating the mechanisms of repression of LIN-28 during larval development—also utilizing whole animal sampling—observed that at later developmental stages, when LIN-28 protein levels were downregulated, LIN-28 mRNA could be detected in association with polyribosome fractions during sucrose gradient analysis, or associated with ribosomes as indicated by ribosome profiling (Seggerson et al. 2002; Stadler et al. 2012). Our finding that the *lep-5* lncRNA plays a significant role in the downregulation of LIN-28 during later larval stages by degrading newly synthesized LIN-28 offers an explanation for the previously observed phenomenon of LIN-28 mRNA translation during these stages of larval development.

*C. elegans* lacking the endogenous *lin-28* 3’ UTR exhibits a superficially wild-type appearance despite the absence of negative regulation by the *let-7* and *lin-4* microRNAs. This contrasts with the *lin-28(ΔCSs)* mutant, which exhibits retarded developmental phenotypes owing to LIN-28 overexpression. This result led us to hypothesize that 3’ UTR sequences outside of the microRNA sites exert a positive effect on LIN-28 expression that can balance the repression mediated by the *lin-4-CS* and *let-7-CS*. The *lin-28* 3’ UTR appears to differ in this respect from the 3’ UTRs of other components of the heterochronic cascade, particularly *lin-14* and *hbl-1*, where deletion of most or all of their respective 3’ UTRs result in pronounced gain-of-function phenotypes that resemble the phenotypes associated with loss-of-function mutations of *lin-4* or *let-7* microRNAs (Wightman et al. 1993; Ilbay and Ambros 2019).

A prior study has demonstrated that mature *let-7* microRNA accumulates to increasing levels as *C. elegans* development progresses (Nahar et al. 2024). Given the long-estimated half-life of *let-7* microRNA, its final concentration at the L4 stage—when *let-7* levels dictate the fates of seam and vulval cells—presumably results from the integration of *let-7* biogenesis and turnover dynamics throughout larval development. Despite the superficially wild-type phenotype of *lin-28(Δ3’ UTR)* animals, the dynamics of LIN-28 protein levels are not normal. We observed lower LIN-28::GFP levels in L1 samples, coupled with elevated levels in L4 samples. The elevated levels of LIN-28::GFP in *lin-28(Δ3’ UTR)* mutants during the L4 stage suggest that *let-7* microRNA production in this mutant may be partially inhibited at later developmental stages. Conversely, the lower levels of LIN-28::GFP at the L1 stage of *lin-28(Δ3’ UTR)* suggest increased *let-7* microRNA production during early larval stages. This implies that the wild-type phenotype of *lin-28(Δ3’ UTR)* mutants may reflect a hyper-accumulation of *let-7* microRNA during early larval stages, which persists into later developmental stages and compensates for the reduced *let-7* production caused by late-stage LIN-28 misexpression.

The individual introductions of three distinct segments of the *lin-28* 3’ UTR resulted in LIN-28 gain-of-function phenotypes, demonstrating that each element is sufficient to promote LIN-28 activity independently of the remainder of the 3’ UTR and that these segments do not necessarily function to counteract the negative regulation imposed by the *let-7-CS* and *lin-4-CS*. Likewise, introduction of the *let-7-CS* and *lin-4-CS* in *lin-28(Δ3’ UTR)* animals was sufficient to suppress the LIN-28 gain-of-function phenotypes observed in the absence of *lep-5* lncRNA thereby indicating that these microRNA binding sites can function independently of the remaining 3’ UTR and their action does not necessarily require counteracting the activities of the positive regulatory elements.

Early L1 *lin-28(Δ3’ UTR)* animals exhibit approximately 25% of the LIN-28 mRNA levels observed in wild-type, suggesting that specific regions within the *lin-28* 3’ UTR contribute to the stabilization of LIN-28 mRNA. Consequently, upon the introduction of one of the LIN-28-promoting 3’ UTR segments, we observed a significant increase in LIN-28 mRNA levels relative to the *lin-28(Δ3’ UTR)* samples. This finding indicates that this element facilitates the stabilization of LIN-28 mRNA in the *lin-28(Δ3’ UTR)* context, leading to LIN-28 gain-of-function. However, the incorporation of this singular region did not fully restore LIN-28 mRNA levels, suggesting that additional regions within the 3’ UTR, likely including the other two blocks of positively acting sequences that we identified, further enhance mRNA stability. The mechanisms through which these 3’ UTR regions stabilize LIN-28 mRNA remain unclear and necessitate further investigation.

In *C. elegans*, the *lin-28* locus produces two LIN-28 isoforms that differ in their N-termini (Seggerson et al. 2002; Moss and Tang 2003). Both N-termini, along with the C-terminus, are predicted to be intrinsically disordered (Varadi et al. 2022; Abramson et al. 2024; UniProt 2025). A notable characteristic of many RNA-binding proteins is the presence of unstructured intrinsically disordered domains (IDRs), which can play crucial roles in the protein’s function (Ottoz and Berchowitz 2020). The observation that GFP-tagging of the LIN-28 C-terminus subtly impairs LIN-28 function suggests that the GFP tag may compromise an essential function of the C-terminal IDR.

The phenotypic similarities observed between *lin-28* gain-of-function alleles and the *let-7* null allele, coupled with the significant negative regulation that LIN-28 exerts on *let-7* microRNA expression, suggest that reduced levels of *let-7* microRNA are a primary factor contributing to the developmental retardation observed in *lin-28* gain-of-function mutants. Notably, the phenotypes exhibited in *lin-28* 3’ UTR mutants that also lacked the *lep-5* lncRNA, were more severe than *let-7(0)* mutants. This observation suggests that the phenotypes manifested in these double mutants are likely attributable to the deregulation of targets of LIN-28 and/or *lep-5* lcnRNA in addition to the *let-7* microRNA.

In *C. elegans*, the maintenance of optimal temporal expression levels of LIN-28 is critical for ensuring robust development and fecundity. In this study, we have demonstrated that *C. elegans* have evolved multiple regulatory mechanisms—both positive and negative—to ensure that LIN-28 is highly expressed during early larval stages and expressed at lower levels during later larval stages, irrespective of environmental conditions. This is particularly evident in *lin-28(Δ3’ UTR)* animals, which exhibit a wild-type phenotype under standard laboratory conditions but display emergent retarded developmental phenotypes resulting from exposure to *P. aeruginosa*, fluctuations in temperature, or the removal of the LIN-28 negative regulator *lep-5* lncRNA. This conditionality of *lin-28* gain-of-function phenotypes suggests a robustness function for the multidimensional, layered regulation of LIN-28 described herein—3’ UTR-dependent regulation of LIN-28 production through the convergence of *lin-4* and *let-7* microRNAs and putative positive factors, as well as 3’ UTR-independent regulation of LIN-28 turnover by the LEP-2*/lep-5* lncRNA system. Further studies are required to elucidate how these modes that regulate LIN-28 may be coupled to physiological and genetic signals, such that LIN-28 developmental dynamics are robustly linked to developmental progression in wild type animals.

## MATERIALS AND METHODS

### Nematode methods and phenotypic analyses

*C. elegans* were cultured on nematode growth medium (NGM) (Brenner 1974) and fed with *E. coli* HB101. Synchronized populations of developmentally staged worms were obtained using standard methods (Stiernagle 2006). A list of strains used in this study is in Table S2.

For heterochronic phenotype analyses, one to four late L4 larvae were picked from healthy uncrowded cultures and placed onto individual NGM plates that were seeded with HB101. Animals were serially transferred daily for three days. Progeny on the plates were left to develop and scored for heterochronic phenotypes as young adults. Unless noted otherwise, all heterochronic phenotype analyses were performed at 15°C. Fluorescence microscopy was used to score COL-19::GFP.

#### Development on Pseudomonas aeruginosa

Development of *C. elegans* on *P. aeruginosa* was performed as previously described (Mirza et al., 2023).

### Microscopy and image quantification

All DIC and fluorescent images were obtained using a W1-Yokogawa spinning disk Nikon confocal microscope equipped with a Prime BSI Express A22E726013 camera, a PLAN APO λD 60x OIL OFN25 DIC N2 objective, and the Nikon Elements software. Prior to imaging, worms were anesthetized with 0.2 mM levamisole in M9 buffer and mounted on 2% agarose pads. All LIN-28::GFP images were taken using the same microscopy settings. Pixel intensity of unmodified images was quantified using Fiji software. Adobe Photoshop was used to adjust the brightness and contrast of representative images used for all figures to enhance the visualization of the DIC and fluorescent signals. Identical brightness and contrast adjustments were used for LIN-28::GFP fluorescent images that are displayed for comparison.

#### Caenorhabditis genomes

All genomes used in this study were provided by the *Caenorhabditis* Genomes Project (http://caenorhabditis.org).

#### Identification of *lin-28* homologs

Identification of *lin-28* homologs was performed using the *Caenorhabditis* Genomes Project BLAST webpage (http://blast.caenorhabditis.org/).

### RNA extraction

Populations of animals were grown at 20°C, collected in M9 buffer, flash-frozen in liquid nitrogen, and total RNA was extracted by adding approximately 20 µL of 100 µm zirconium beads (OPS Diagnostics) and homogenized in a Next Advance Bullet Blender Blue at setting 8 for 2 minutes at 4°C with Qiazol reagent (Qiagen). The remainder of the RNA extraction protocol is as described by (McJunkin and Ambros 2017).

### Protein extraction

Populations of animals were grown at 20°C, collected in M9 buffer, and flash-frozen in liquid nitrogen. Protein was extracted by adding an equal volume of 2x Laemmli buffer without dye (20% glycerol, 4% SDS, 0.125M Tris [pH 6.8], and 0.1 M DTT) and approximately 20 µL of 100 µm zirconium beads (OPS Diagnostics), homogenized in a Next Advance Bullet Blender Blue at setting 8 for 2 minutes at 4°C, heated for 5 minutes at 99°C, centrifuged at 16.1 rcf for 5 minutes at room temperature, and the supernatant was transferred to a fresh tube.

### Western blotting

Western blotting was performed as described previously (Dutta and Baehrecke 2008). Primary antibodies used were rabbit anti-LIN-28 (1:10,000) (Seggerson et al. 2002) and mouse anti-alpha-tubulin (1:10,000) (Sigma T6074). Western blots were imaged using Amersham Imager 600.

It is important to note that our western blot analyses utilizing the anti-LIN-28 antibody displayed bands that migrated inconsistently with the expected molecular weight of LIN-28 isoforms; none of these bands exhibited a shift in size in samples containing LIN-28::GFP, indicating that they are cross-reacting peptides unrelated to LIN-28. To further validate this observation, we employed CRISPR/Cas9 to delete the entire coding sequence of *lin-28*, resulting in the loss of the two bands presumed to correspond to LIN-28A and LIN-28B, while the suspected non-specific bands remained (*lin-28(ma637)*; Fig. S1d).

### Quantitative PCR

For all mRNA quantification, cDNA was synthesized using SuperScript IV (Invitrogen) following the manufacturer’s instructions. For all small RNA quantification, cDNA was synthesized using miRNA 1^st^ Strand cDNA Synthesis Kit (by stem-loop) (Vazyme) following the manufacturer’s instructions. qPCR reactions were performed using 2x UltraSYBR Mixture (Low ROX) (CoWin Biosciences) following the manufacturer’s instructions, using a QuantStudio 12K Flex (Applied Biosystems). For all LIN-28 mRNA experiments, ΔCTs were calculated by normalizing samples to ACT-1/2 mRNA. For all *let-7* microRNA experiments, ΔCTs were calculated by normalizing samples to U18 snoRNA. All oligos used for cDNA synthesis and qPCR in this study are described in Table S3.

### Transgenic constructs

The extrachromosomal array maEx268, which was used to rescue *lep-5* in the strains VT4303 and VT4307, was generated by PCR cloning the *lep-5* locus. The resulting PCR product was column purified and injected along with the *rol-6(su1006)*-containing plasmid pRF-4 into *lep-5(ma613)* mutants. The extrachromosomal array maEx269, which was used to rescue *lep-5* in the strains VT4400, VT4411, VT4412 and VT4413, was generated by PCR cloning the *lep-5* locus, and by PCR cloning *rol-6(su1006)* from the plasmid pRF-4 and *Pmyo-2::mCherry* from the plasmid pCFJ90. The resulting PCR products were column purified and injected into *lep-5(ma613)* mutants. All oligos used to generate transgenic constructs in this study are described in Table S3.

### Targeted genome editing by CRISPR/Cas9

Mutants were generated using CRISPR/Cas9 methods adapted from (Nelson and Ambros 2019; Nelson and Ambros 2021). See Table S4 for a detailed list of alleles and Table S3 for descriptions of the oligos used in allele generation in this study. All alleles listed with two single-stranded oligodeoxynucleotides (ssODNs) as HR templates utilized two ssODNs with overlapping complementarity and singled stranded 5’ overhangs in the generation of the allele.

## ACKNOWLEDGEMENTS

We thank the members of the Ambros and the Mello laboratories for helpful discussions and comments on this project. We thank Christopher Hammell for sending us the LIN-28 antibody.

## AUTHOR CONTRIBUTIONS

Conceptualization: C.N., V.R.A.; Methodology: C.N., V.R.A.; Formal analysis: C.N., V.R.A.; Investigation: C.N., V.R.A.; Resources: V.R.A.; Data curation: C.N.; Writing - original draft: C.N.; Writing - review & editing: C.N., V.R.A.; Supervision: V.R.A.; Project administration: V.R.A.; Funding acquisition: V.R.A.

