## Supplemental figures for "Multi-dimensional regulation of LIN-28 temporal expression dynamics in the *C. elegans* heterochronic gene cascade"

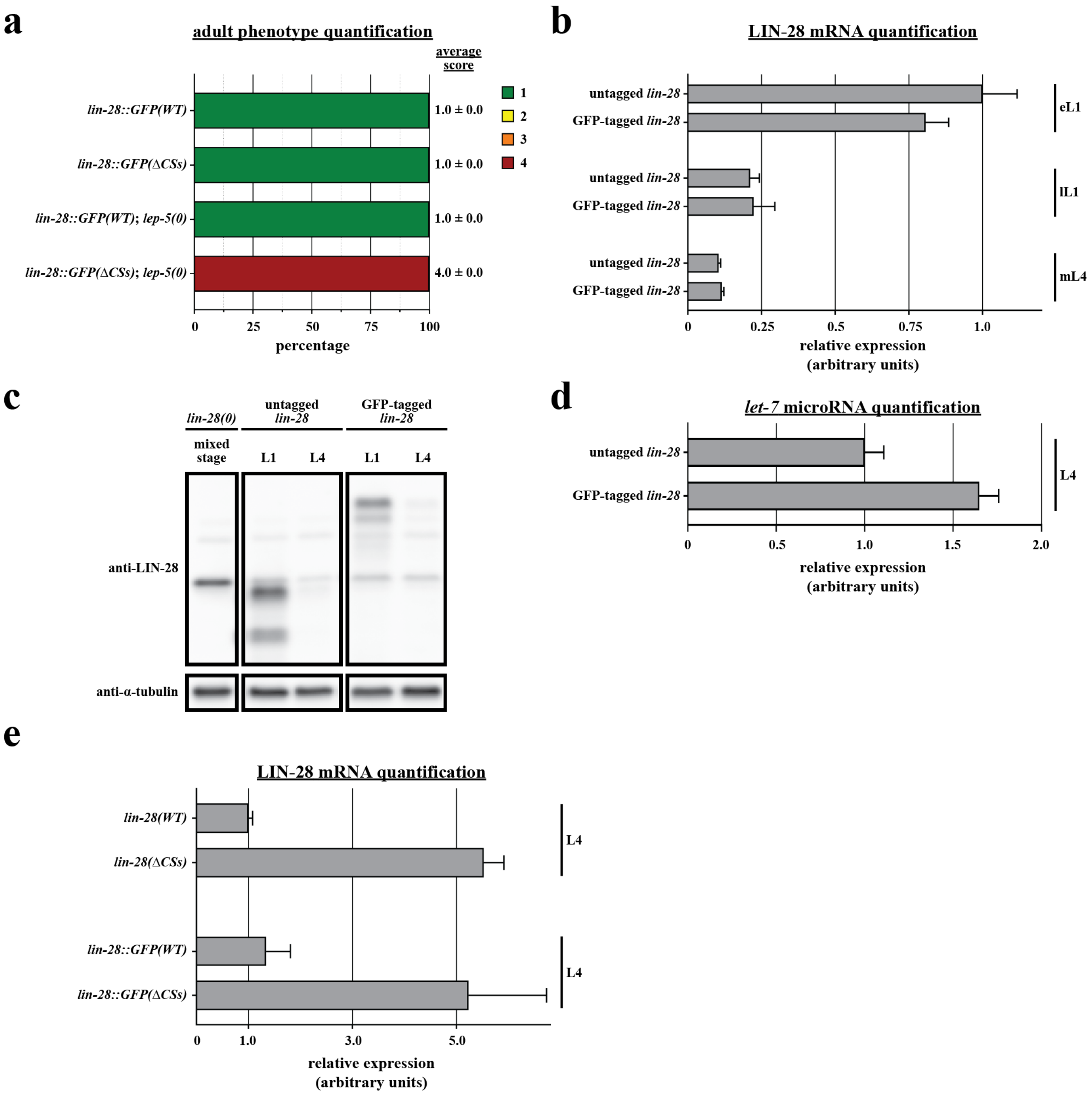


**Figure S1. GFP tagging impairs the function of LIN-28.**

(a) Quantification of adult phenotypes of *lin-28::GFP* animals with either a wild-type 3' UTR or ΔCSs in the presence and absence of *lep-5*. N's (from top to bottom) = 52, 99, 74, and 51. Data are presented as mean score ± standard deviation.

(b) RT-qPCR analysis of LIN-28 mRNA in whole animal extracts from early L1, late L1, and mid L4 stages of wild-type untagged and GFP-tagged *lin-28* strains. N = 4. Data are presented as mean ± standard deviation.

(c) Whole animal lysates from *lin-28* null animals (mixed stage sample), untagged LIN-28 (L1 and L4 samples), and GFP-tagged LIN-28 (L1 and L4 samples) were analyzed using western blotting with an anti-LIN-28 antibody.

(d) RT-qPCR analysis of *let-7* microRNA levels in mid L4 stage whole animal extracts of wild-type untagged and GFP-tagged *lin-28* strains. N = 5. Data are presented as mean ± standard deviation.

(e) RT-qPCR analysis of LIN-28 mRNA in whole animal extracts from L4 wild-type untagged and GFP-tagged *lin-28* strains. N = 3. Data are presented as mean ± standard deviation.

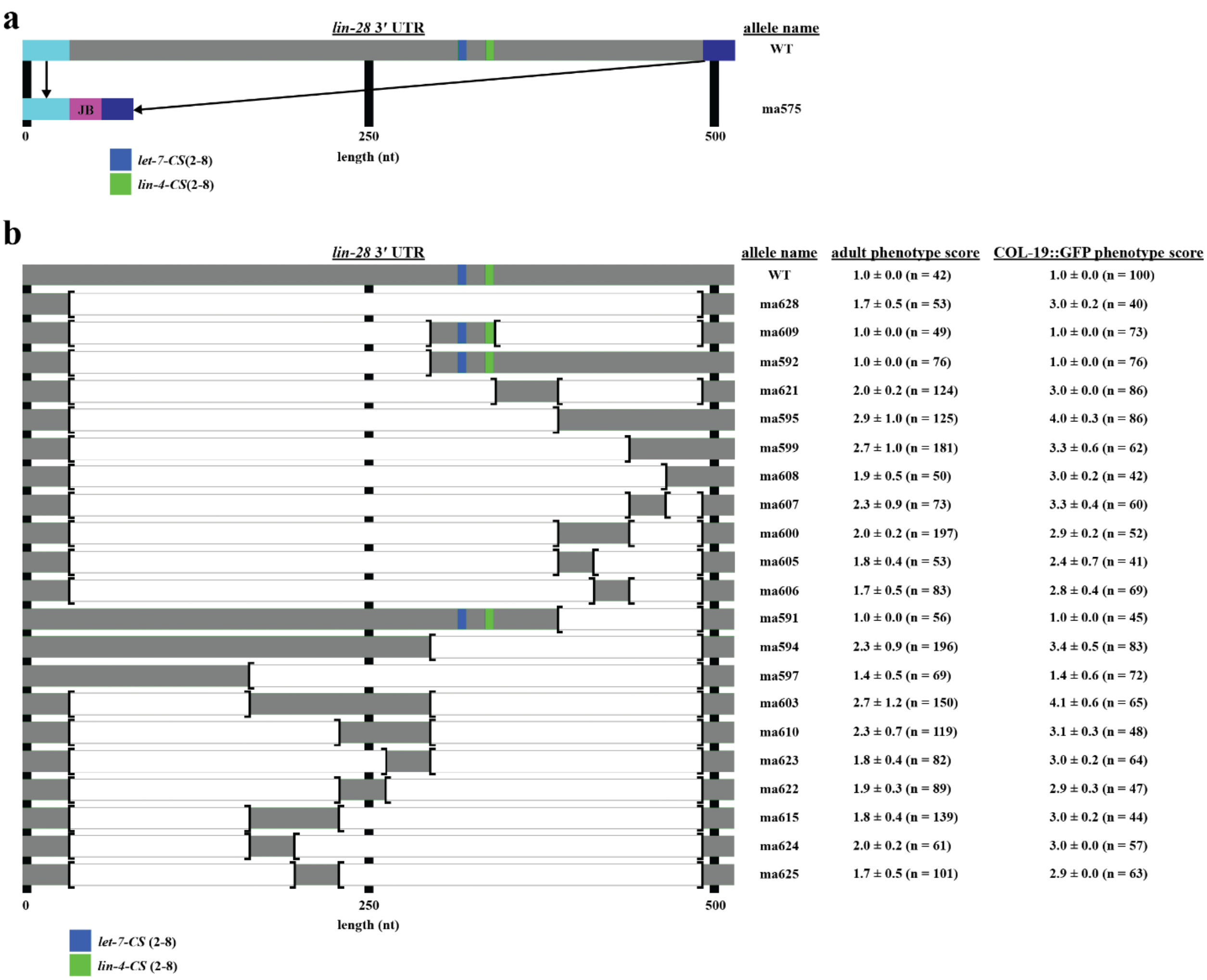


**Figure S2. 3' UTR truncations of *lin-28* generated to identify positive regulatory elements.**

(a) Depiction of the wild-type *C. elegans lin-28* 3' UTR and *lin-28(ma575)*, which replaces 458 bp of the endogenous *lin-28* 3' UTR with the "jump-board" (JB) sequence (artificial CRISPR landing site; magenta) (Duan et al. 2020).

(b) Depiction of the endogenous *lin-28* 3' UTR truncation strains generated to isolate regions that contain positive regulatory elements and quantifications of adult and COL-19::GFP phenotype scores. Data are presented as mean score ± standard deviation.
